# Integrating gene annotation with orthology inference at scale

**DOI:** 10.1101/2022.09.08.507143

**Authors:** Bogdan M. Kirilenko, Chetan Munegowda, Ekaterina Osipova, David Jebb, Virag Sharma, Moritz Blumer, Ariadna E. Morales, Alexis-Walid Ahmed, Dimitrios-Georgios Kontopoulos, Leon Hilgers, Kerstin Lindblad-Toh, Elinor K. Karlsson, Zoonomia Consortium, Michael Hiller

## Abstract

Annotating coding genes and inferring orthologs are two classical challenges in genomics and evolutionary biology that have traditionally been approached separately, limiting scalability. We present TOGA, a method that integrates structural gene annotation and orthology inference. TOGA implements a different paradigm to infer orthologous loci, improves ortholog detection and annotation of conserved genes compared to state-of-the-art methods, and handles even highly-fragmented assemblies. TOGA scales to hundreds of genomes, which we demonstrate by applying it to 488 placental mammal and 501 bird assemblies, creating the largest comparative gene resources so far. Additionally, TOGA detects gene losses, enables selection screens, and automatically provides a superior measure of mammalian genome quality. Together, TOGA is a powerful and scalable method to annotate and compare genes in the genomic era.

## One-Sentence Summary

A scalable gene annotation approach using a different paradigm to detect orthologous loci provides comparative data for hundreds of genomes.

Homologous genes have a common evolutionary ancestry. Orthologs are homologous genes that originated from a speciation event, whereas paralogs originated from a duplication event. Distinguishing orthologs and paralogs is a fundamental problem in evolutionary and molecular biology (*1*) and a prerequisite for many genomic analyses, including reconstructing phylogenetic trees, predicting gene function, investigating molecular and genome evolution, and discovering differences in genes that underlie phenotypes of the sequenced species (*2–6*).

Current methods for orthology inference are either based on graph or gene tree approaches or a combination of both (*7*). Graph-based methods cluster genes into pairs or groups of orthologs based on pairwise sequence similarity such as (reciprocal) best alignment hits (*8–12*). Gene tree-based methods determine whether the evolutionary lineages of two genes coalesce in a speciation or a duplication node (*12–14*). Importantly, these approaches analyze coding or protein sequences of genes, necessitating the identification of gene locations (structural gene annotation) in each genome before inferring orthologs. This has two limitations. First, gene annotation quality has a large influence on the accuracy of orthology inference (*15*). Second, generating high-quality annotations is time-consuming and typically requires comprehensive transcriptomics (gene expression) data, leading to a growing gap between genome sequencing and annotation, including orthology inference.

Here, we developed TOGA (Tool to infer Orthologs from Genome Alignments), an integrative pipeline that jointly addresses two fundamental problems in genomics and evolutionary biology: structural gene annotation and orthology inference.

## Results

### A different paradigm for orthology detection

All orthology detection methods implicitly or explicitly use the principle that orthologous sequences are generally more similar to each other than to paralogous sequences (*1*). While existing methods focus on similarity between coding sequences that typically evolve under purifying selection, this principle also extends to non-exonic regions (introns, intergenic regions) that largely evolve neutrally. The key innovation implemented in TOGA is that intronic and flanking intergenic regions of orthologous gene loci are also more similar to each other, provided that the evolutionary distance between two species is short enough to retain sequence similarity in neutrally evolving regions. For example, the evolutionary distance between human and other placental mammals and between chicken and other birds is ≤0.55 substitutions per neutral site (fig. S1, Tables S1-S2), explaining why orthologous introns and intergenic regions partially align within these clades (Fig. 1A,E and fig. S2). In contrast, evolutionary distances between paralogs that duplicated before the divergence of these clades often exceed 1 substitution per neutral site, resulting in unaligned introns and intergenic regions. TOGA exploits this principle by (i) taking a well annotated genome such as human, mouse or chicken as a reference, (ii) inferring all (co-)orthologous gene loci from a genome alignment between reference and a query species (other placental mammals or birds), and (iii) annotating and classifying these genes (Fig. 1B-D).

**Fig. 1:**
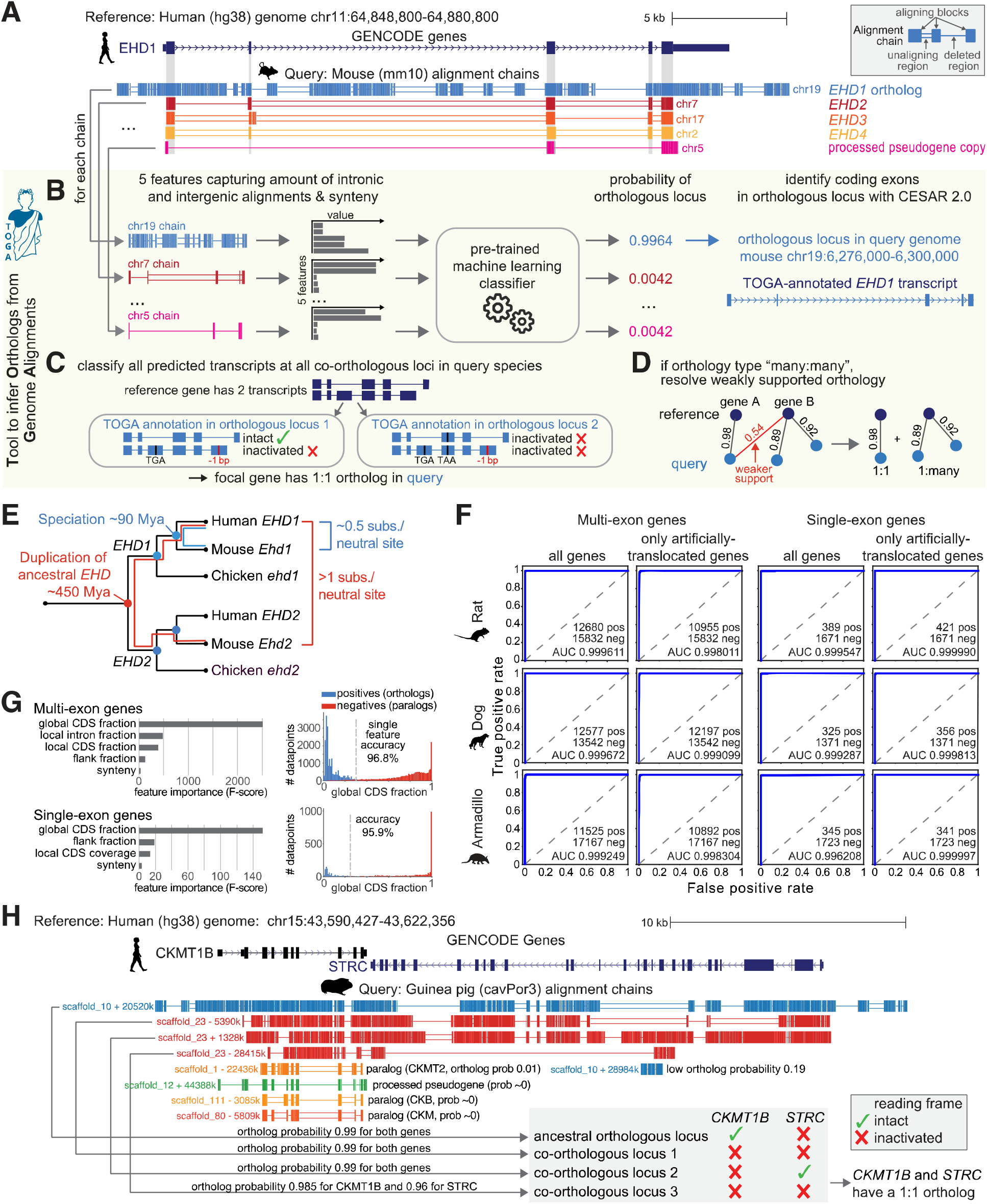
TOGA utilizes intronic and intergenic alignments to detect orthologous gene loci. (A) UCSC genome browser view of the human *EHD1* gene locus shows five alignment chains to mouse. Only the orthologous chr19 locus but not paralogous (chr7/17/2) and processed pseudogene (chr5) loci show intronic and intergenic alignments. (B-D) Illustration of the TOGA pipeline steps that identify orthologous loci, annotate and classify transcripts, and resolve weak orthology connections. (E) Evolutionary distance explains why only the orthologous *EHD1* locus shows intronic and intergenic alignments. (F) Orthology detection performance shown as Receiver Operating Characteristics curves for single- and multi-exon genes as well as for genes that lack synteny due to deliberately-introduced translocations. (G) Feature importance for detecting orthologous genes and the distribution of the most important feature (“global CDS fraction”; proportion of coding exon alignments of all aligning chain blocks). (H) Importance of detecting all orthologous loci and determining reading frame intactness. The human *STRC* and *CKMT1B* locus is quadruplicated in guinea pig (top four chains). TOGA correctly recognizes all four co-orthologous loci. Despite the quadruplication, TOGA finds that only one copy of each gene encodes an intact reading frame and correctly infers a 1:1 orthology relationship.

### The TOGA annotation and orthology detection pipeline

TOGA takes as input a gene annotation of the reference and a whole genome alignment between reference and query genome. TOGA infers orthologous loci in the query, annotates genes, determines orthology types (number of orthologs per gene in reference and query as 1:1, 1:many, many:1, or many:many), detects lost genes, and generates protein/codon alignments. In the first step, TOGA uses a pairwise genome alignment between reference and query, represented by chains of co-linear local alignments (*16*). These alignment chains capture both orthologous gene loci as well as loci containing paralogs or processed pseudogenes (Fig. 1A). To distinguish between them, TOGA computes characteristic features that capture the amount of intronic and intergenic alignments, considering each gene and each overlapping chain (Fig. 1B and fig. S3). Synteny (conserved gene order), which can help to distinguish orthologs from paralogs (*14*), is used as an additional feature. TOGA then uses machine learning to compute the probability that a chain represents an orthologous locus for the gene of interest.

To train the machine learning classifier, we used known orthologous genes between human (reference) and mouse (query) from Ensembl Compara (*14*) (fig. S4). Testing this classifier on independent query species (rat, dog, armadillo) that represent different placental mammalian orders showed a near perfect classification of orthologous chains (Fig. 1F, Table S3). Manual investigation of misclassifications showed that false positives mostly represent partial or full gene duplications (actual co-orthologous loci) and that half of the false negatives may be related to faster X chromosome evolution (*17*) (fig. S5-S6). Features capturing intronic/intergenic alignments are most important for the classification performance (Fig. 1G). In contrast, synteny is the least important feature, likely reflecting our training data sets that we deliberately enriched with translocated orthologs (fig. S7). Using synteny as an auxiliary but not determining feature enables TOGA to also accurately detect orthologs that underwent translocations or inversions (fig. S8).

In a second step, for every transcript of a reference gene, TOGA uses CESAR 2.0 (*18, 19*) to determine the positions of coding exons of the focal gene in each (co-)orthologous query locus (Fig. 1B and fig. S9-S10). Since orthologous gene loci do not necessarily encode a gene with an intact reading frame (Fig. 1H), TOGA assesses reading frame intactness for each transcript (Fig. 1C and fig. S11). To this end, TOGA implements an improved version of our gene loss detection approach (*5*) and identifies gene-inactivating mutations (frameshifting, stop codon or splice site mutations, exon or gene deletions) while taking assembly incompleteness into account (fig. S12-S17). A gene is only classified as lost, if all transcripts at all (co-)orthologous loci are classified as lost. TOGA detects gene losses using the mutations present in the assembly without attempting to fix potential base errors (fig. S18-S19). We benchmarked the specificity of this approach on 11,161 conserved genes. Only 21, 22, 12 and 21 of these genes are misclassified as inactivated in mouse, rat, cow and dog, respectively, indicating a very high specificity of 99.80-99.89% (Table S4). Manual inspection showed that misclassified cases include highly-diverged genes, genes that evolved drastic changes in exon-intron structure or protein length, and a lost gene that is compensated by a processed pseudogene copy, which highlights cases of less certain gene conservation (fig. S20-S23).

In the third step, TOGA determines the orthology type by considering all reference genes and all orthologous query loci that encode an intact reading frame (Fig. 1C and fig. S24). Finally, TOGA uses an orthology graph approach to resolve weakly-supported orthology relationships among many:many orthologs (Fig. 1D and fig. S25).

### TOGA improves ortholog detection

To assess the performance of TOGA’s orthology detection pipeline, we compared it against Ensembl Compara, which integrates graph- and tree-based methods (*14*). Using orthologs between human and three representative mammals (rat, cow, elephant), TOGA detected 97.6%, 98.9% and 96.5% of the orthologs provided by Ensembl (Fig. 2A, Table S5), showing a good agreement. Furthermore, for >90% of these commonly-detected orthologs, TOGA inferred the same orthology type (Fig. 2B). A quarter of the discrepancies are cases where TOGA infers 1:1 and Ensembl 1:many. In several of these cases, Ensembl annotates a processed pseudogene copy as a second ortholog (fig. S26).

**Fig. 2:**
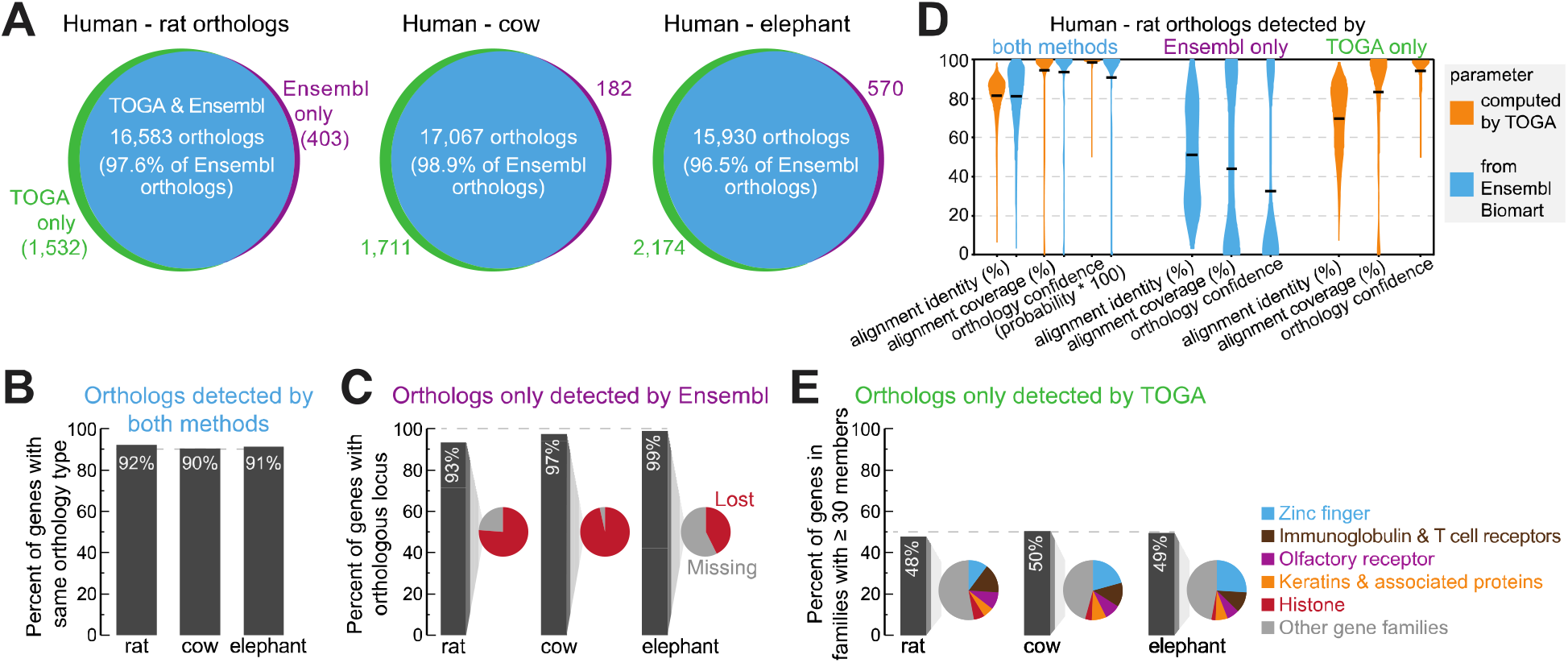
TOGA improves ortholog detection. (A) Ortholog overlap between Ensembl Compara and TOGA. (B) Percent of commonly-detected orthologs having the same orthology type. (C) Percent of orthologs only detected by Ensembl, for which TOGA detects an orthologous locus but classifies the gene as lost or missing. (D) Human-rat orthologs detected by both or only one method. Violin plots compare identity and coverage of coding region alignments and orthology confidence probabilities. Note that for orthologs only detected by TOGA, these features are not available on Ensembl Biomart, and vice versa. Horizontal black lines represent the mean. (E) Percent of orthologs only detected by TOGA that belong to gene families with ≥30 members. Pie charts show the proportion of the most frequent gene families.

For the orthologs detected only by Ensembl, TOGA did identify an orthologous locus in >93% of the cases, but detected either reading frame inactivating mutations, indicating a lost gene, or that more than half of the coding region overlaps assembly gaps in the query (classified as a missing gene) (Fig. 2C and fig. S27-S28). Consistent with these cases including more questionable orthologs, parameters measuring alignment identity (mean 51%), alignment coverage (mean 44%) and orthology confidence (mean 32%) are substantially lower compared to orthologs detected by both methods (means 81%, 94%, 91%) (Fig. 2D).

TOGA predicted for the three species 1,532 (rat), 1,711 (cow) and 2,174 (elephant) additional orthologs that are not listed in Ensembl (Fig. 2A). For rat, this includes *PAX1*, an important developmental transcription factor that was potentially missed by Ensembl because of a mis-annotated N-terminus (fig. S29). About half of these genes belong to large families such as zinc fingers, olfactory receptors or keratin-associated proteins (Fig. 2E). These genes exhibit alignment identity (mean 70%), alignment coverage (mean 83%) and orthology confidence (mean 94%) values that are more similar to orthologs detected by both methods (means 82%, 94%, 99%) (Fig. 2D), supporting that these genes are undetected orthologs.

### TOGA improves annotation of conserved genes

We performed a direct comparison between TOGA’s comparative gene annotations and annotations generated by Ensembl and by the NCBI Eukaryotic Genome Annotation Pipeline (*20, 21*), two state-of-the-art methods that integrate transcriptomics data, homology-based data and *ab initio* gene predictions. We first applied TOGA using the human GENCODE 38 annotation (*22*) as the reference to other placental mammals that have Ensembl (70 species) or NCBI (118 species) annotations. We then used BUSCO (Benchmarking Universal Single-Copy Orthologs; odb10 dataset) to compare the percent of completely detected near-universally-conserved mammalian genes (*23*). TOGA annotations have a higher completeness score for 97% (Ensembl) and 91.5% (NCBI) of the species (Fig. 3A,B, Tables S6-S7), increasing annotation completeness of conserved genes by an average of 4.1% or ∼377 genes (Ensembl) and 0.7% or ∼64 genes (NCBI) (fig. S30).

**Fig. 3:**
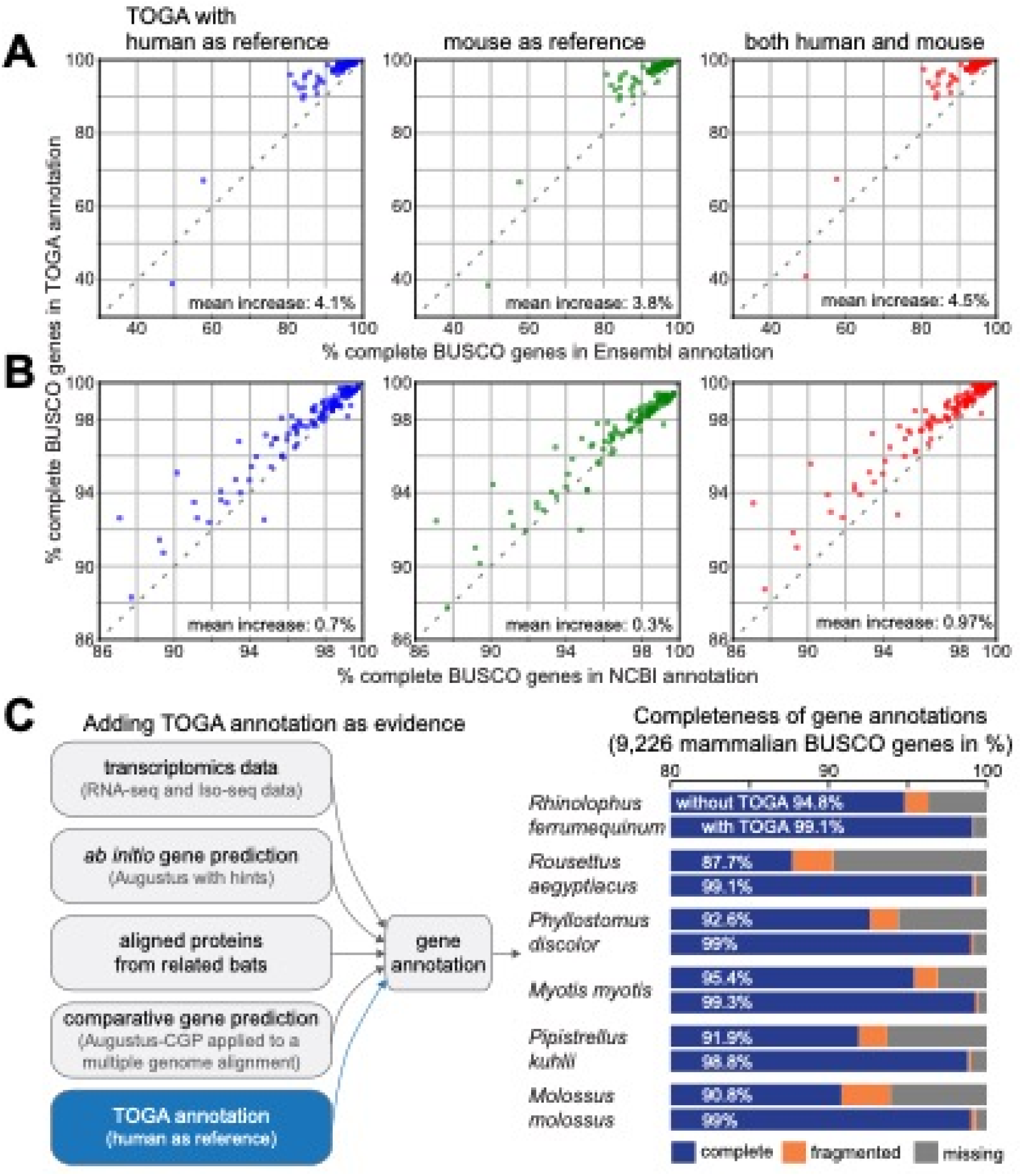
TOGA improves annotation of conserved genes. (A,B) Completeness of mammalian BUSCO genes in annotations generated by TOGA (Y-axis), Ensembl (X-axis in A) and the NCBI Eukaryotic Genome Annotation Pipeline (X-axis in B). Each dot represents one species. The set of 70 and 118 species in A and B overlaps but is not identical. (C) Gene evidence used to annotate six bat species. Adding TOGA as evidence increases annotation completeness of mammalian BUSCO genes by 3.9% to 11.4%.

Second, we used TOGA with the mouse GENCODE M25 annotation (*22*) as the reference. This resulted in a higher BUSCO completeness for 98.5% (Ensembl) and 64% (NCBI) of the species (Fig. 3A,B, Tables S6-S7). As a homology-based method, TOGA benefits from the quality and comprehensiveness of the human and mouse input annotation (*21, 22*). However, homology-based methods cannot annotate orthologs of genes that are not present in the reference (fig. S31, Table S8). This downside can be counteracted by combining multiple references. Indeed, combining human- and mouse-based TOGA annotations achieves a higher BUSCO completeness for almost all species (>98%) (Fig. 3A,B).

Third, further adding TOGA annotations with generated additional references (cow, horse and cat) increases the total number of annotated genes and detects additional lineage-restricted genes (fig. S32-S34). Nevertheless, comprehensive annotation of lineage-specific exons and genes requires transcriptomics data or *ab initio* predictions (fig. S35).

### TOGA improves annotations even if transcriptomics data are available

Transcriptomics data provides direct evidence of transcripts expressed in the sampled tissues. We next tested whether TOGA can improve annotation of conserved genes, even if transcriptomics data and other gene evidence are available. To this end, we used six high-quality bat genomes (*6*) and first annotated genes by integrating available transcriptomics data, *ab initio* gene predictions (Augustus (*24*)), aligned proteins from closely related bats, and comparative gene predictions (Augustus-CGP applied to a multiple genome alignment (*25*)). For the six bats, these annotations contained 87.7% to 95.4% of the genes in the mammalian BUSCO odb10 set (Fig. 3C, Table S9). Adding TOGA with human as the reference generated annotations containing 98.8% to 99.3% of the BUSCO genes. This shows that even if comprehensive gene evidence are available, TOGA can improve the annotation of conserved genes.

### TOGA joins split genes in fragmented assemblies

Genes split between different scaffolds are often missed or annotated as fragments, hampering downstream analyses. Although current genome projects aim to generate highly-complete, chromosome-level assemblies (*6, 26*), even such assemblies can contain fragmented genes (fig. S36). Furthermore, many currently available mammal or bird assemblies exhibit fragmentation (*27, 28*). To improve comparative annotation and orthology inference of fragmented genes, we leveraged TOGA’s ability to detect orthologous loci of gene fragments. We implemented a gene joining procedure that recognizes orthologous parts of 1:1 orthologous genes, joins them together, and generates an annotation and codon/protein alignments for the full gene (Fig. 4A and fig. S37).

**Fig. 4:**
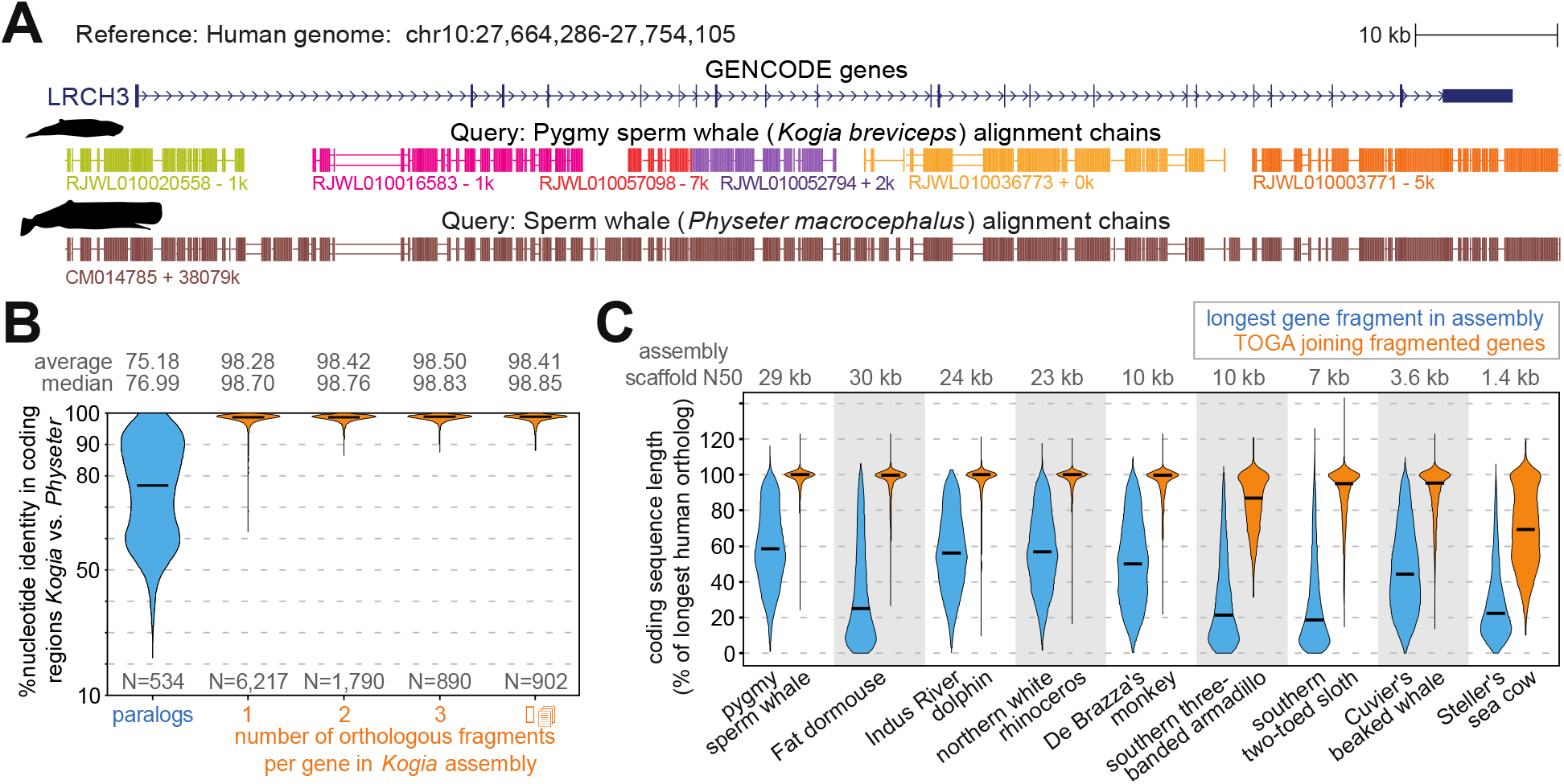
TOGA accurately joins genes split in fragmented genome assemblies. (A) The ortholog of human *LRCH3* is split into six fragments (evident by six chains) in the highly-fragmented pygmy sperm whale (*Kogia breviceps*) assembly (*27*). Different chain colors represent different scaffolds. TOGA correctly detects and joins all six orthologous gene fragments. The highly-contiguous assembly of the closely related sperm whale (*Physeter macrocephalus*) (*29*), where *LRCH3* is located on a single scaffold, shows a highly-similar alignment block structure. (B) Violin plots show the coding exon identity between *Kogia breviceps* and *Physeter macrocephalus*. Horizontal black lines represent the median. Fragmented orthologs joined by TOGA have an identity distribution highly-similar to orthologs already present on a single scaffold. (C) Violin plots compare the coding sequence length before (blue) and after joining split genes (orange). Length is relative to the longest transcript of the human ortholog. Codon insertions can increase the relative length to >100%.

To evaluate the accuracy of this procedure, we leveraged that sequences of orthologous but not paralogous genes from closely related species are expected to be highly similar. Indeed, comparing a highly-fragmented with a highly-contiguous assembly of two sperm whale species (*27, 29*) showed that orthologous genes located on a single scaffold in both species have a much higher sequence identity (mean 98.70%) than paralogous genes (mean 75.18%) (Fig. 4B). Consequently, if TOGA would misidentify paralogous fragments as orthologs, sequence identity should decrease for fragmented genes. However, we observed an equally high identity for orthologous genes joined from two, three or even more fragments (Fig. 4B), indicating a high accuracy.

Demonstrating the effectiveness of TOGA’s gene joining procedure, in the highly-fragmented sperm whale assembly the mean coding sequence length after joining fragmented genes is 97% of the length of the orthologous human gene. This is a substantial improvement over the single largest orthologous fragment present in the assembly (mean 59%) (Fig. 4C, Table S10). We obtained similar improvements for other highly-fragmented assemblies. Even for an assembly of the extinct Steller’s sea cow with a scaffold N50 value of just 1.4 kb (*30*), TOGA improved the relative coding sequence length from 28% to 70%. Thus, TOGA increases the utility of fragmented genomes for comparative analyses.

### TOGA scales to hundreds of genomes

As complete genomes are generated at an increasing rate, annotation and orthology inference methods that can handle hundreds or thousands of genomes are needed. Unlike previous methods, TOGA’s reference-based methodology scales linearly with the number of query species. We leveraged this by applying TOGA with the human GENCODE 38 annotation (19,464 genes) as reference to a large set of placental mammals, comprising 488 assemblies of 427 distinct species (Fig. 5A, Tables S1,S11). As expected, TOGA annotates more orthologous genes in the six Hominoidea (ape) species that are closely related to human (median 19,192). Importantly, for the remaining 482 assemblies, TOGA also annotated a median of 18,049 orthologs, indicating that TOGA is an effective annotation method across placental mammals.

**Fig. 5:**
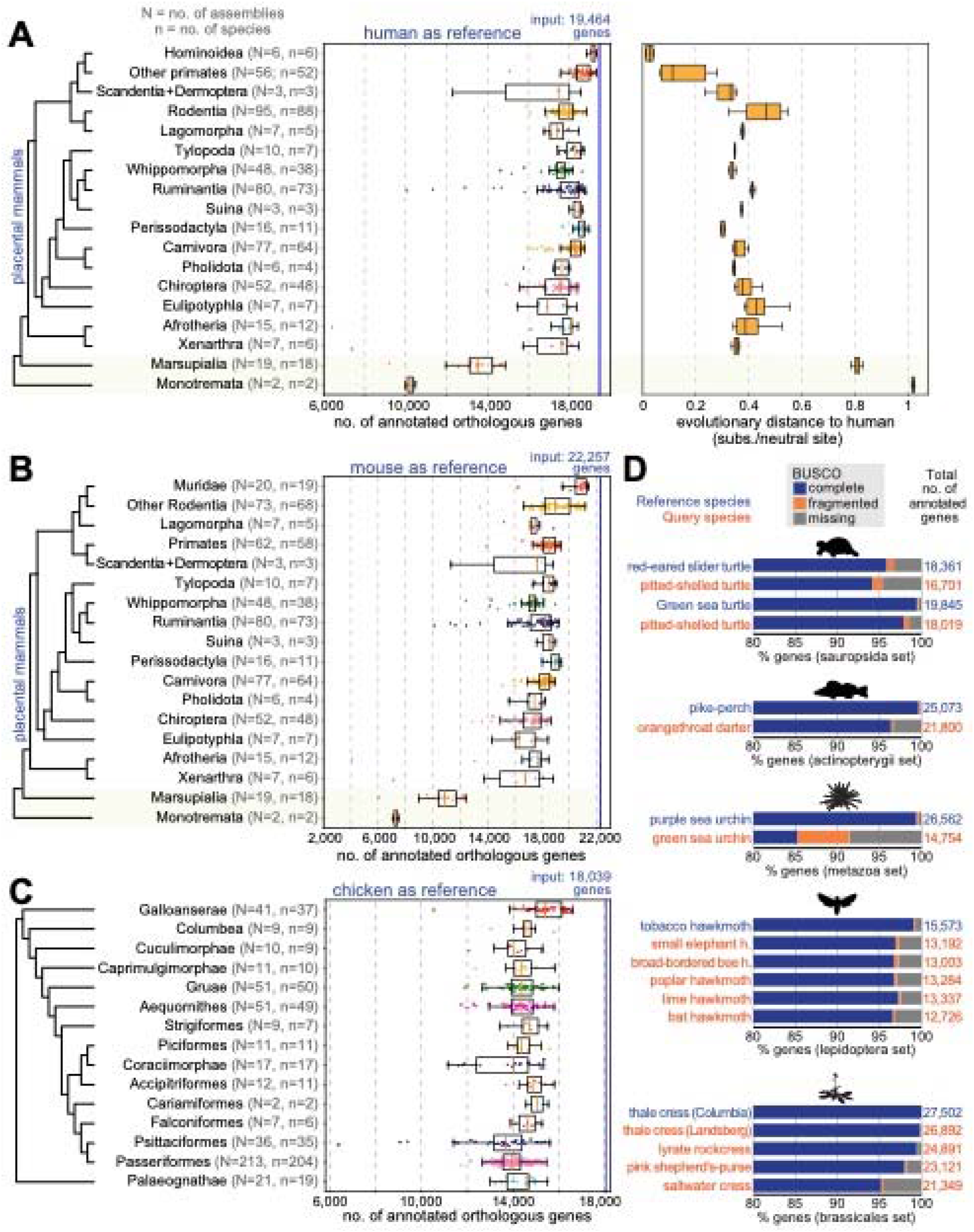
Large-scale application of TOGA to hundreds of genomes. (A) Human as reference. Left: Box plots with overlaid data points show the number of annotated orthologs. Non-placental mammals are highlighted with a yellow background. Right: Box plots showing evolutionary distances to human. (B) Mouse as the reference. Muridae are shown as a separate group. (C) TOGA with chicken as the reference, applied to 501 bird assemblies. (D) TOGA for other species using NCBI RefSeq annotations (*21*) as the reference. BUSCO gene completeness of the reference annotation provides an upper bound for the completeness of TOGA’s query annotation.

Fitting generalized linear models shows that the number of annotated orthologs is positively correlated with assembly quality metrics (contig and scaffold N50) and negatively correlated with the evolutionary distance (substitutions per neutral site) and divergence time (millions of years) to human (fig. S38, Table S12). Evolutionary distance has a stronger influence than divergence time. This is exemplified for Perissodactyla, where TOGA consistently annotates more orthologs than in many rodents, despite the rodent lineage splitting from human more recently.

To explore the influence of the reference genome, we applied TOGA to the same 488 placental mammal assemblies using the mouse GENCODE M25 annotation (22,257 genes) as reference (Fig. 5B, Table S1). Corroborating a general influence of evolutionary distance and divergence time, TOGA annotated more orthologs for the 20 closely related Muridae assemblies (median 20,918) than for the remaining 466 assemblies (median 18,115). Overall, the number of annotated genes is similar to the human-based annotations.

### TOGA provides a superior approach for assessing mammalian assembly quality

TOGA’s gene classification also provides a powerful benchmark to measure assembly completeness and quality. To this end, we first compiled a comprehensive set of 18,430 ancestral placental mammal genes, defined as human coding genes that have an intact reading frame in the basal placental clades Afrotheria and Xenarthra (Table S13). For each of the 488 assemblies, we then used TOGA’s gene classification to determine which percent of these ancestral genes have an intact reading frame without missing sequence. This completeness measure is significantly correlated with the completeness value computed by BUSCO in genome mode (Pearson r = 0.73, P=10^-81^) (Fig. 6A). However, BUSCO’s values saturate at ∼97% for highly complete assemblies, while TOGA’s completeness values exhibit a larger dynamic range (Fig. 6A,B), providing a better resolution to distinguish highly contiguous from less contiguous assemblies. This is exemplified by two closely related bats: a high-quality *Rhinolophus ferrumequinum* and a less-contiguous *R. sinicus* assembly have similar BUSCO (96.4% vs. 96.3% complete genes) but different TOGA completeness values (94.4% vs. 88.2%) (Fig. 6C). These results are driven by the TOGA methodology and not by the twofold increased gene number (18,430 vs. 9,226 genes; fig. S39).

**Fig. 6:**
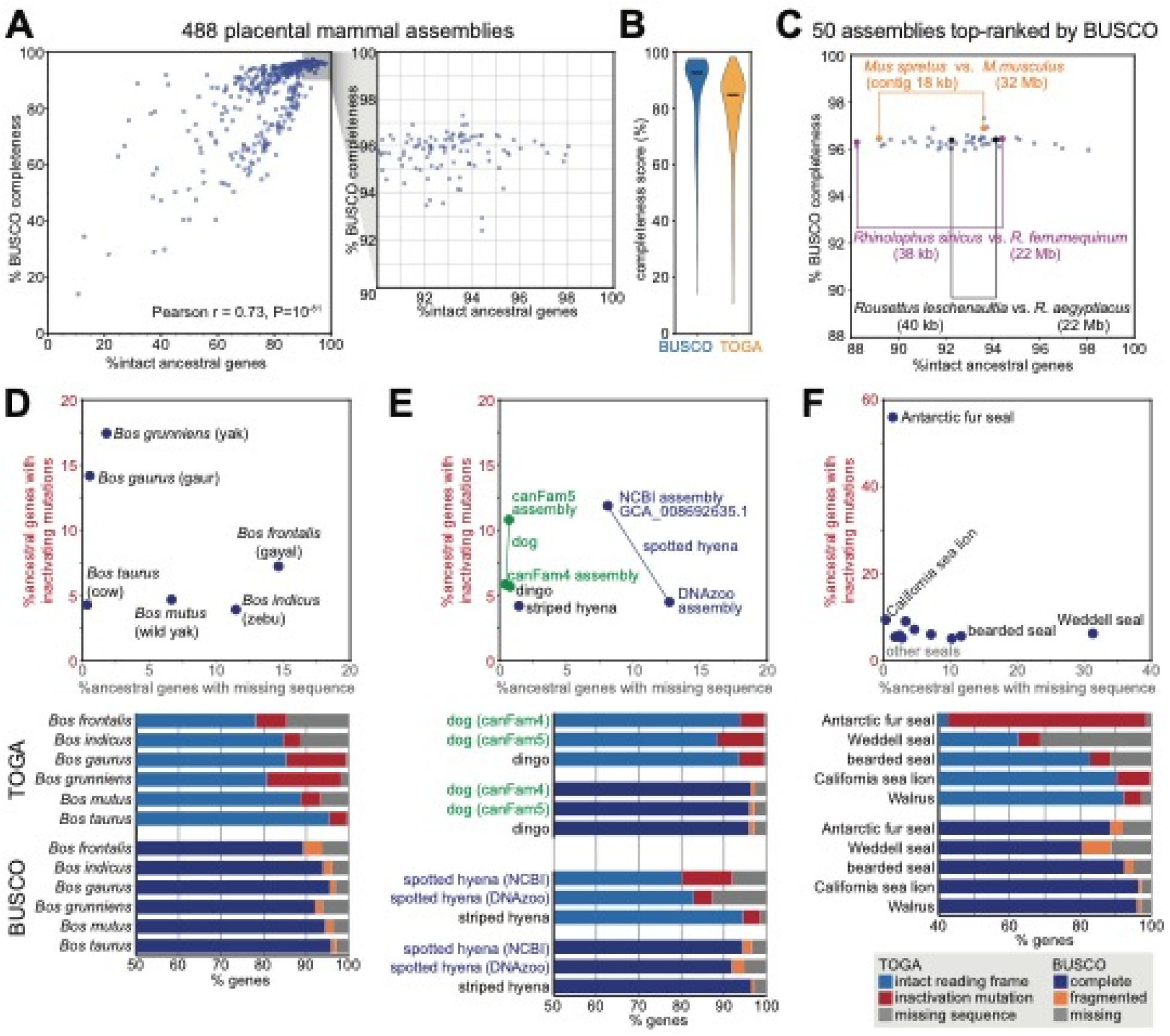
TOGA provides a superior measure of mammalian assembly quality. (A) Comparison of the percent complete BUSCO genes and TOGA’s percent intact ancestral genes for 488 placental mammal assemblies. Each dot represents one assembly. (B) Violin plots of BUSCO’s and TOGA’s completeness values. Horizontal black lines represent the median. (C) BUSCO’s and TOGA’s completeness values for 50 assemblies that are top-ranked by BUSCO. Three pairs of closely related species are highlighted that have different assembly contiguity (contig N50) values and are distinguishable in terms of gene completeness by TOGA, but not by BUSCO. (D-F) TOGA distinguishes between genes with missing sequences and genes with inactivating mutations. This highlights assemblies with a higher incompleteness or base error rate that is often not detectable by the BUSCO metrics.

BUSCO’s fragmented or missing gene classification indicates how much of the gene was detected, but does not distinguish between the two major underlying reasons: assembly incompleteness that results in missing gene sequence vs. assembly base errors that destroy the reading frame. TOGA’s gene classification explicitly distinguishes between these two different assembly issues, which provides valuable information on assembly quality. For example, TOGA detects a higher percentage of genes exhibiting inactivating mutations in the *Bos gaurus* (gaur, 14.2%) compared to the *Bos taurus* (cow, 4.3%) assembly, indicating that the *B. gaurus* assembly has an elevated base error rate, whereas both assemblies are indistinguishable in terms of BUSCO completeness (95.8 vs. 95.5%) (Fig. 6D). Similarly, the dog canFam5 assembly exhibits an elevated base error rate compared to dog canFam4 or dingo, whereas all three assemblies have similar BUSCO scores (Fig. 6E). Assemblies of the same species can suffer from different issues, illustrated by the spotted hyena, where the NCBI GCA_008692635.1 assembly has less missing sequence, but a noticeably higher base error rate compared to the DNAzoo assembly (Fig. 6E). Finally, illustrating extreme cases among seals, 56% of the genes in the Antarctic fur seal have inactivating mutations and 31% of the genes in the Weddell seal have missing exonic sequence (Fig. 6F).

### TOGA facilitates more accurate codon alignments

Codon or protein alignments are important to screen for selection patterns and reconstruct phylogenetic trees, but alignment errors can substantially impact the outcome (*31*). TOGA implements two features that help to avoid codon alignment errors. First, TOGA masks all gene-inactivating mutations such as frameshifts, which otherwise can result in misalignments (fig. S40). Second, whereas existing methods align entire orthologous coding sequences, TOGA is aware of orthology at the exon level. This enables an “exon-by-exon” procedure that generates alignments by aligning and joining individual orthologous exons, which can avoid alignment errors (fig. S41).

### Applying TOGA to 501 bird as well as other non-mammalian genomes

To demonstrate TOGA’s applicability to non-mammalian genomes, we used chicken (18,039 genes, RefSeq annotation (*21*)) as the reference and applied TOGA with default models and parameters to 501 assemblies of 476 distinct bird species (*28, 32*) (Tables S11,S14). Across all assemblies, TOGA annotated a median of 14,058 orthologous genes (Fig. 5C, Table S14).

We also explored whether TOGA can be applied to species other than mammals and birds. Tests with turtles, fish, sea urchins, hawk moths and Brassicaceae plants provide encouraging results (Fig. 5D) that may be further improved by retraining the machine learning classifier, defining new features, and adjusting genome alignment parameters and CESAR’s splice site profiles.

### Comprehensive resources for comparative genomics

For the 488 placental mammal and 501 bird assemblies, we provide comparative gene annotations, ortholog sets, lists of inactivated genes and multiple codon alignments generated with MACSE v2 (*33*) for download at http://genome.senckenberg.de/download/TOGA/. To our knowledge, these comprise the largest comparative genomics datasets for both clades so far. To facilitate visualizing and analyzing these data, we implemented a TOGA annotation track for the UCSC genome browser (*34*) (fig. S42). Our UCSC browser mirror at https://genome.senckenberg.de/ provides these annotation tracks for all analyzed assemblies.

## Discussion

We envision two main use cases of TOGA. First, by detecting inactivated genes and providing orthologous sequences for codon alignments, TOGA enables phylogenomic analyses as well as screens for selection patterns and gene losses that are linked to relevant phenotypes (*6, 35–38*). Second, TOGA can provide an initial annotation of conserved genes for newly-sequenced genomes or be integrated together with available transcriptomics data and *ab initio* gene predictions to comprehensively annotate conserved and lineage-specific genes. Additionally, TOGA’s classification of ancestral genes provides a useful assembly quality benchmark.

TOGA’s application range comprises species with “alignable” genomes, which we define in our context as genomes where orthologous neutrally evolving regions partially align. In general, this holds for evolutionary distances of ∼0.6 substitutions per neutral site. Interestingly, applying TOGA with human as the reference to 18 marsupial and two monotreme mammals, where neutrally evolving regions are diverged because of the larger evolutionary distance (∼0.8 and ∼1 substitution per neutral site between human and marsupials/monotremes), still annotates on average 13,397 and 10,238 orthologs (Fig. 5A,B), primarily because gene order is conserved (fig. S43). Nevertheless, for these more distant clades, human is not a powerful reference and a marsupial and a monotreme mammal should be used as the reference instead.

With the tree of life becoming more densely populated with genomes, thanks to great efforts of large-scale projects and numerous laboratories (*26–28, 39*), TOGA provides a general strategy to cope with the annotation and orthology inference bottleneck. For every “alignable” clade of interest, one can select one, or ideally several, reference species. Assembly and annotation of the reference(s) should ideally be highly complete, and reference choice can be influenced by the evolutionary distance to focal query species. References can be defined for different taxonomic ranks, from the class to the family or genus level. For example, in the Bat1K project (*40*), we aim at generating a high-quality assembly and comprehensive gene annotation for representatives of all bat families to serve as references for dozens or hundreds of other bats in these families.

## Materials and Methods

### TOGA input and output

As input, TOGA requires (i) the reference and query genome file in 2bit format (an indexed and compressed file that can be generated from a multi-fasta file with UCSC genome browser tool twoBitToFa), (ii) the coding gene annotation of the reference genome in bed-12 format (can be generated from genePred or gtf formats with the UCSC utilities genePredToBed and gtfToGenePred), and (iii) a chain file containing chains of co-linear local alignments between the reference and query genome. Optionally, information about U12 introns, where non-canonical splice sites are common, can be provided as input. If the gene annotation provides more than one transcript for a gene, TOGA will process all transcripts, as detailed below. To generate high-quality annotations, we recommend including representative isoforms (sometimes called principal) for each gene, in particular those that capture differences in exon-intron structures, but to exclude isoforms that represent much shorter and likely non-functional transcripts such as potential targets for nonsense-mediated decay. We also recommend excluding transcripts that represent fusion isoforms between two ancestral genes, as including such fusion transcripts interferes with inferring the correct orthology type.

TOGA provides rich output and generates (i) a gene annotation of the query species in bed-12 format, (ii) an annotation file listing processed pseudogenes detected in the query in bed-9 format, (iii) the protein and codon alignments of all annotated genes in fasta format, (iv) per-exon nucleotide alignments together with alignment quality scores (nucleotide and protein similarity) in fasta format, (v) a table listing orthology relationships between genes in the reference and query (orthology type as 1:1, 1:many, etc.), (vi) a table of genes, transcripts and projections that are classified as intact, lost or other states describing the likelihood that a functional protein is encoded, (vii) a list of all detected gene-inactivating mutations in tsv format, (viii) a table listing for each reference transcript which alignment chains overlap this transcript and what their ortholog score is, (ix) tab-separated files that can be loaded as UCSC genome browser tracks to visualize the annotations, chain classification scores, exon-intron structure with inactivating mutations, and exon and protein alignments with nucleotide identity and BLOSUM alignment scores.

### Overview of TOGA

The pipeline implemented in TOGA consists of the following steps. First, for each coding gene annotated in the reference, TOGA applies machine learning to determine orthologous (and co-orthologous) loci in the query genome by inferring which alignment chains represent orthologous alignments. Second, for each (co-)orthologous locus in the query genome, TOGA uses CESAR 2.0 (*18, 19*) to determine the positions and boundaries of all coding exons of each gene. In this step, TOGA also analyses the reading frame of the annotated transcript, filters the resulting exon alignments, detects gene-inactivating mutations, determines whether undetected exons are missing due to assembly gaps, and classifies the annotated transcript as intact, missing or inactivated. Third, after inferring all orthologous loci and annotating all genes, TOGA infers the orthology type between genes and resolves spurious many:many relationships that are only supported by weak orthology. The three steps are described in detail in the following.

### Inferring orthologous loci from pairwise genome alignments

In the first step, TOGA infers orthologous loci by using pairwise chains of co-linear local alignments, computed between a reference and query genome (see below), and the gene annotation of the reference genome.

#### Identifying candidate chains

TOGA first extracts all chains that overlap or span at least one coding exon for a given coding gene. Since a naive approach that loops over all possible gene-chain pairs is time-consuming, TOGA implements a faster approach that relies on sorting genes and chains. Specifically, for each chromosome or scaffold, TOGA sorts the genomic regions of all genes and all chains by the start coordinate in the reference genome. Then, for each chain, TOGA iterates over the sorted list of genes, starting with the first gene that intersected the previous chain (all upstream genes can be skipped). For each gene, we determine whether the chain overlaps or spans at least one coding exon, which makes this chain a candidate chain. The iteration is stopped at the first gene that starts downstream of the current chain end. Compared to the naive approach, this procedure also has an asymptotic quadratic runtime of O(N^2^), but only in the worst case where every chain overlaps every gene. In practice, we found that this procedure results in a speedup of ∼60 fold (human vs. mouse, 0.5 vs. 30 min), because it avoids considering numerous genes that are upstream or downstream of a focal chain.

#### Feature extraction for machine learning

Given a gene and an overlapping chain, TOGA computes the following features by intersecting the reference coordinates of aligning blocks in the chain with different gene parts (coding exons, UTR (untranslated region) exons, introns) and the respective intergenic regions. We define the following variables (see also fig. S3).

- *c*: number of reference bases in the intersection between chain blocks and coding exons of the gene under consideration.
- C: number of reference bases in the intersection between chain blocks and coding exons of all genes.
- a: number of reference bases in the intersection between chain blocks and coding exons and introns of the gene under consideration.
- A: number of reference bases in the intersection between chain blocks and coding exons and introns of all genes and the intersection between chain blocks and intergenic regions (excludes UTRs).
- f: number of reference bases in chain blocks overlapping the 10 kb flanks of the gene under consideration. Alignment blocks overlapping exons of another gene that is located in these 10 kb flanks are ignored.
- i: number of reference bases in the intersection between chain blocks and introns of the gene under consideration.
- CDS (coding sequence): length of the coding region of the gene under consideration.
- I: sum of all intron lengths of the gene under consideration.

Using these variables, TOGA computes the following features:

- “global CDS fraction” as C / A. Chains with a high value have alignments that largely overlap coding exons, which is a hallmark of paralogous or processed pseudogene chains. In contrast, chains with a low value also align many intronic and intergenic regions, which is a hallmark of orthologous chains.
- “local CDS fraction” as c / a. Orthologous chains tend to have a lower value, as intronic regions partially align. This feature is not computed for single-exon genes.
- “local intron fraction” as i / I. Orthologous chains tend to have a higher value. This feature is not computed for single-exon genes.
- “flank fraction” as f / 20,000. Orthologous chains tend to have higher values, as flanking intergenic regions partially align. This feature is important to detect orthologous loci of single-exon genes.
- “synteny” as log10 of the number of genes, whose coding exons overlap by at least one base aligning blocks of this chain. Orthologous chains tend to cover several genes located in a conserved order, resulting in higher synteny values, which can help to distinguish orthologs from paralogs (14, 41–43).
- “local CDS coverage” as c / CDS, which is only used for single-exon genes.

The term ‘global’ refers here to features computed from all genes that overlap the chain, whereas ‘local’ refers to features computed from just the single gene under consideration. Most of these features quantify how well intronic and intergenic regions, which largely evolve neutrally, align in comparison to coding exons, which largely evolve under purifying selection. Because selection in UTR exons is variable, alignments overlapping UTR exons are ignored for feature computation. All features are visually explained in fig. S3.

#### Generating training data of orthologous and non-orthologous genes

We trained a machine learning approach to use the above-described features to distinguish chains representing alignments to orthologous genes. As training data, we used human-mouse 1:1 orthologs from Ensembl (*44*) (Release 97, downloaded July 2019), for which the “orthology confidence” feature is 1. For each gene, we only considered the transcript with the longest coding region.

As positives (orthologous chains), we selected those chain-gene pairs, where (i) the chain is the top-level (highest-scoring) chain covering the gene, and (ii) the chain represents a true orthologous alignment of the gene (fig. S4). The latter condition was implemented by requiring that the Ensembl-annotated mouse ortholog is located at the query coordinates provided by this chain. To obtain negatives (non-orthologous chains that typically represent alignments to paralogs or processed pseudogenes), we reasoned that by definition other chains overlapping exons of true 1:1 orthologous genes cannot represent co-orthologs. Consequently, such chains represent non-orthologous alignments and were added to the negative set (fig. S4). To avoid selecting negative chains that cover only a small fraction of the gene, we only considered non-orthologous chains, where aligning blocks overlap at least 35% of coding exons. Furthermore, for the positive and negative sets, we only considered chains with a score of at least 7,500 and genes whose coding exons overlap less than 75 different chains.

We noticed that the vast majority of positives had high synteny feature values, indicating that inversions or translocations, which break the co-linear order between genes, are rare among human-mouse 1:1 orthologs. Since we aimed at also accurately detecting orthologous genes that underwent genomic rearrangements, we enriched the positive training dataset with artificially rearranged chain-gene pairs, generated by trimming long syntenic chains to new single gene-covering chains. To this end, we considered all 1:1 orthologous genes whose orthologous chain is among the top 100 scoring orthologous chains already used in the positive training set. For each of these genes, we determined breakpoints of an artificial rearrangement by adding a random number ranging from -10,000 to 3,000 to the gene start (transcription start) and adding a random number ranging from -3,000 to 10,000 to the gene end (transcription end). As a result, the artificial rearrangement may even lack some parts of the beginning or end of the gene (fig. S7). However, to avoid cases where the artificial rearrangement lacks most of the coding exons, we only considered artificial rearrangements that include at least 80% of the gene’s coding region. For each artificial rearrangement, we used the breakpoints to trim the original orthologous chain, resulting in a new chain that typically covers only a single gene and sometimes only a part of a single gene (fig. S7).

To create the final training dataset with balanced proportions, we combined all 14,376 real orthologous and all 5,844 artificially rearranged gene-chain pairs as the positive set (20,220 entries) and considered 20,220 randomly chosen gene-chain pairs as the negative set. We then split this training dataset into single- and multi-exon genes to train the two models, as described below. To create independent test datasets, we applied the same procedure to genome alignments of different query species, human-to-rat, human-to-dog and human-to-armadillo.

#### Model training and testing

We trained two separate models (one for multi-exon genes and one for single-exon genes), since two features that quantify intronic alignments (“local CDS fraction” and “local intron fraction”) can only be computed for multi-exon genes. For single-exon genes, we found the feature “local CDS coverage” to be helpful in detecting orthologous loci. We did not use this feature when training the multi-exon model, as it did not increase classification performance further and hampered the detection of partial lineage-specific duplications of multi-exon genes. Hence, the multi-exon model was trained using all six features except “local CDS coverage” and the single-exon model was trained using all six features except “local CDS fraction” and “local intron fraction” (fig. S3).

We used the XGBoost (*45*) gradient boosting library, a machine learning approach that was successfully applied to a variety of classification tasks, to train both models with the following parameters: number of trees: 50, maximal tree depth: 3, learning rate: 0.1. For each gene-chain pair, the XGBoost predictor outputs a score between [0,1] that the chain represents an orthologous locus for the gene. The single-exon gene model showed a 5-fold cross-validation accuracy of 99.41% (standard deviation 0.28%) and the multi-exon gene model showed a 5-fold cross-validation accuracy of 99.23% (standard deviation 0.07%).

To assess the importance of the features for chain classification (Fig. 1G), we computed the “gain” value (*45*), which measures the contribution of the feature for each decision tree in the gradient boosting model as the average reduction of the loss function that is obtained when using this feature for splitting the training data.

We tested the single- and multi-exon model on independent test sets obtained for three representative placental mammals that include both a close sister species to mouse (rat) and more distant outgroups (dog, armadillo). To evaluate the performance of the models in detecting translocated or inverted orthologous genes, we separately tested them on real orthologous genes (typically high synteny values) and artificially rearranged orthologous genes (typically low synteny values of log_10_(1)) (Fig. 1F, Table S3). ROC curves were computed by ranking each gene-chain pair by the orthology score.

#### Chain classification

To annotate genes and infer orthologs, we consider in this manuscript all gene-chain pairs where the orthology score is ≥0.5. This threshold can be adjusted by users via a TOGA parameter.

#### Annotating processed pseudogenes

Chains also align processed pseudogene copies of multi-exon genes, which enables TOGA to augment the query genome annotation by annotating processed pseudogenes. To this end, TOGA implements a post-hoc classification of non-orthologous chains into those that represent paralogs vs. processed pseudogene copies. To distinguish between paralogous and processed pseudogenes, TOGA computes for multi-exon genes the “alignment to query span” value. Defining *e* as the number of reference bases in the intersection between chain blocks and exons (here using both UTR and CDS) and defining *Q* as the span of the chain in the query genome, “alignment to query span” is computed as *e* / *Q*. This value is close to 1 for chains representing alignments to processed pseudogenes in the query, as introns are completely ‘deleted’ and thus the summed length of exon alignments is similar to the chain length in the query. Non-orthologous chains where the “alignment to query span” value is > 0.95 and that overlap only one gene are classified as processed pseudogene chains. TOGA then uses the chain span to annotate the processed pseudogene copy in the query and correctly label this locus as such (fig. S44).

#### Gene-spanning chains

For genes that are entirely absent from the query genome, either because they are deleted or completely overlap assembly gaps, there can be a chain that spans this gene but none of its aligning blocks overlap exons of this gene. Since the machine learning step cannot be applied to these chains, as most features cannot be computed, TOGA treats these chains as follows, but only if the focal gene completely lacks a detected orthologous locus. If aligning blocks of this chain overlap coding exons of at least two other genes, we consider it as an orthologous chain candidate for the focal gene. For such chains, TOGA runs CESAR 2.0 on the query locus defined by the closest upstream and downstream aligning blocks, if the distance is at most 1 Mb or at most 50 times the gene length (CDS start – CDS end). CESAR may detect the gene or remnants of it in this query locus, even if the gene did not align at the nucleotide level in the genome alignment chains. As described below, TOGA then filters the CESAR output to determine whether the gene exists but was missed in the genome alignment chain, whether the gene is likely deleted, or whether the gene overlaps assembly gaps and is thus missing.

### Transcript alignment and classification

#### CESAR alignment

The result of the first step is a set of gene-chain pairs that are classified as orthologous and provide an orthologous locus for the respective gene in the query genome. In the second step, TOGA identifies the loci and splice site boundaries of all coding exons by aligning the coding exons of the reference species to the query locus. To this end, TOGA individually considers all transcripts provided for this gene and uses CESAR (Codon Exon Structure Aware Realigner) version 2.0 (*18, 19*) in multi-exon mode. Briefly, CESAR 2.0 is a Hidden Markov model-based method that takes the coding exons of the reference species as input and considers reading frame and splice site information when generating exon alignments in the query sequence. CESAR has a high accuracy in correctly aligning shifted splice sites, is able to detect precise intron deletions that merge two neighboring exons, and generates alignments of intact exons (defined as exon alignments with consensus splice sites and an intact reading frame) whenever possible (*18, 19*). Before running CESAR, TOGA replaces in-frame TGA stop codons in the reference sequence, which can encode a selenocysteine amino acid, by NNN. This replacement enables CESAR to align such TGA stop codons to sense or to stop codons. Also, if information about U12 introns in the reference is provided as input, TOGA passes this information to CESAR. As U12 intron splice sites can comprise a variety of dinucleotides including AT-AC, GT-AG, GT-GG, AT-AT or AT-AA (*46*), we have changed the U12 donor and acceptor splice site profile in CESAR to capture this splice site diversity with a uniform nucleotide distribution. Since knowledge about U12 introns in the reference may be incomplete or not always available, TOGA considers every intron in the reference without canonical GT/GC-AG splice sites as a putative U12 intron. For human or mouse as the reference, we used U12 data from U12DB (*47*).

#### Exon classification

After parsing the CESAR output, TOGA classifies each exon as present (P), missing (M) or deleted (D). This step is necessary as the Viterbi algorithm used in CESAR’s HMM may also output alignments of exons that do not exist in the query locus, either because the exon is truly deleted or diverged to an extent that no meaningful alignment is possible (class D) or because the exon overlaps an assembly gap in the query genome (class M).

To distinguish between classes P, M, and D, TOGA utilizes that an orthologous chain provides not only the orthologous query locus, but the aligning blocks of the chain also provide information about the location of individual exons. TOGA determines whether the CESAR-detected exon location overlaps the query genome locus that should contain the exon according to the genome alignment chain. If this is the case, then both the nucleotide-based genome alignment chain and the codon-based CESAR alignment agree on the exon location in the query, and TOGA classifies these exons as present (P). For exons where the chain and CESAR disagree on the location and for exons that align only with the more sensitive CESAR method, TOGA uses two metrics to evaluate whether the exon aligns better than randomized exons. The first metric is the %nucleotide identity, defined as the percentage of identical bases in the CESAR alignment. The second metric, %BLOSUM, measures the amino acid similarity between reference and query using the BLOSUM62 matrix. Let S_RQ_ be the sum of BLOSUM scores for each amino acid pair between reference (R) and query (Q), with codon insertions and deletions getting a score of -1. As S_RQ_ depends on the length of the exon, we also determine the maximum score possible for this exon by comparing the reference sequence to itself, thus computing S_RR_. %BLOSUM is defined as S_RQ_ / S_RR_ * 100. To determine thresholds that separate real and randomized exon alignments, we extracted 137,935 exons of human-mouse 1:1 orthologous genes for which the TOGA-annotated exon overlaps an Ensembl-annotated exon (real exons). Randomized exon alignments were obtained by aligning real exon sequences to the reversed query sequence with CESAR. By comparing %nucleotide identity and %BLOSUM between real and random CESAR exon alignments, we defined thresholds as %nucleotide identity ≥ 45% and %BLOSUM ≥ 20% (fig. S9). These thresholds correspond to a sensitivity of 0.98 and a precision of 0.99. Exons that exceed these thresholds are classified as present (P). For all other exons, TOGA determines whether the query locus expected to contain this exon overlaps an assembly gap (≥10 consecutive N characters) in the query genome. If so, the exon is classified as missing (M), otherwise it is classified as class D. Exons not spanned by an orthologous chain are also classified as missing (M), as such cases are often due to assembly fragmentation and incompleteness. The exon classification workflow is detailed in fig. S10.

#### Transcript annotation and classification

To annotate transcripts in the query genome, TOGA uses the splice site coordinates of the CESAR alignment to annotate all exons of the given reference transcript that were classified as present in the previous step.

Gene orthology must be inferred based on the number of (co-)orthologs in the query that likely encode a functional protein. For example, even if TOGA detects a single orthologous locus for the given gene with high confidence, the predicted gene could be lost in the query, resulting in a 1:0 orthology relationship (no ortholog in the query). Similarly, as exemplified in Fig. 1H, TOGA can detect four orthologous loci in the query, but if only one of these loci encodes a functional gene, this results in a 1:1 orthology relationship. For these reasons, TOGA implements a transcript classification step to determine whether an annotated transcript is likely or unlikely to encode a functional protein.

Transcript classification is not a straightforward problem, as assembly gaps result in missing parts of the CDS and individual exons can get lost in otherwise clearly conserved genes, as shown in previous work (*5*). To take this complexity into account, we decided to classify annotated transcripts into five different major categories:

- “intact” transcripts, for which the middle 80% of the CDS is present (not missing sequence) and exhibits no gene-inactivating mutation, are likely to encode functional proteins,
- “partially intact” transcripts, for which ≥50% of the CDS is present and the middle 80% of the CDS exhibits no inactivating mutation, may also encode functional proteins, but the evidence is weaker as more of the CDS is missing due to assembly gaps,
- “missing” transcripts, for which less than 50% of the CDS is present and the middle 80% of the CDS exhibits no inactivating mutation, are undecided as more than half of the CDS is missing but no strong evidence for loss exists,
- “uncertain loss” transcripts exhibit at least one inactivating mutation in the middle 80% of the CDS, but evidence is not strong enough to classify the transcript as lost; hence, it may or may not encode a functional protein,
- “lost” transcripts, for which evidence for loss is sufficiently strong, are unlikely to encode a functional protein.

As shown in the flowchart in fig. S11A, TOGA derives this classification by first determining whether the transcript exhibits no (intact, partially intact, missing) or at least one (uncertain loss, lost) gene-inactivating mutation in the middle 80% of the CDS. This key distinction is motivated by our observation that frameshift and stop codon mutations in conserved genes mostly occur in the first or last 10% of the CDS (fig. S12). Fig. S11B illustrates several examples of these five transcript types.

A special and rarely-occurring category called “paralogous projection” refers to cases where no orthologous chain but only a paralogous-classified chain was detected. This can arise if the real orthologous gene is entirely missing in the assembly (thus only a paralog aligns) or if TOGA misclassifies the orthologous gene because of excessive divergence of intronic/intergenic regions. If the locus represented by the paralogous chain does not receive any annotation via an orthologous chain, then TOGA also annotates a gene at this locus (shown in fig. S6), as this locus likely encodes a gene. However, the annotation is labeled as a paralogous projection and shown in brown color.

#### Gene-inactivating mutations

To distinguish between intact and lost transcripts, TOGA considers the following gene-inactivating mutations: frameshifting insertions and deletions, in-frame (premature) stop codons, mutations that disrupt the canonical donor (GT/GC) or acceptor (AG) splice site dinucleotides, and deletions of single or multiple consecutive exons that together are not divisible by three and thus result in a frameshift. Contrary to our previous work (*5*), we do not consider larger frame-preserving deletions as inactivating mutations anymore, because we observed a number of cases where large deletions did occur in otherwise conserved genes. Examples of insertions or deletions (ranging from several hundred to a few thousand base pairs) inside large exons are shown in fig. S16. Examples of deletions of entire exon(s), sometimes comprising seven consecutive exons, are shown in fig. S17. These large frame-preserving deletions result in substantially shorter but likely functional proteins (though it is not known whether the function is truly conserved). Importantly, TOGA does consider stop codons as inactivating that may be assembled at a new exon-exon boundary, which arose from deletions of in-frame exons (fig. S14).

In case of precise intron deletions that merge two neighboring exons into a single larger exon, we do not consider the deletion of the splice sites. For U12 splice sites (labeled as such in the reference or inferred from non-canonical reference splice site dinucleotides), we do not consider splice site mutations. In-frame stop codons that were already present in the reference sequence (selenocysteine-encoding TGA codons or stop codon readthrough) are ignored. Two or more frameshifts that compensate each other (e.g. a -1 and -2 bp deletion, or three -1 bp deletions) and do not result in a stop codon in the new reading frame are not considered as inactivating mutations (fig. S15).

#### Transcript loss criteria

Using the list of detected inactivating mutations, TOGA quantifies the maximum percent of the reading frame that remains intact in the query (fig. S13). To distinguish between “intact”, “partially intact” and “missing” transcripts, we ignore missing sequence (NNN codons) in this calculation. To distinguish between “uncertain loss” and “lost” transcripts, we count missing sequence as aligning codons, making the conservative assumption that missing codons correspond to sense codons in the unknown query sequence (fig. S13), since this procedure results in a consistent classification of transcripts that have the same inactivating mutations and only differ in the amount of missing sequence.

Based on the observation that inactivating mutations in conserved genes rarely occur in the middle 80% of the CDS (fig. S12), transcripts classified as “uncertain loss” or “lost” transcripts exhibit at least one inactivating mutation in the middle 80% of the CDS. The following criteria distinguish between “lost” and “uncertain loss” transcripts. Lost transcripts have a maximum percent intact reading frame <60% and exhibit inactivating mutations in at least 2 coding exons (fig. S11B). The latter requirement is motivated by previous observations that mutations in a single exon of an otherwise-conserved gene are not sufficient to infer gene loss (*5*). For genes with >10 exons, we replace the requirement of mutations in at least two coding exons by requiring mutations in at least 20% of the coding exons. For single exon genes, we simply require two inactivating mutations. As the size of individual exons can be large, we make an exception for multi-exon transcripts, where a single large exon represents a substantial part (≥40%) of the CDS. Such transcripts are also classified as “lost” if at least 2 mutations occurred in this large exon (fig. S11B). All other transcripts that have ≥1 inactivating mutation in the middle 80% of the CDS are classified as “uncertain loss”, indicating that evidence for loss is not strong enough as a larger part of the CDS remains potentially intact (>60%) or not enough exons exhibit inactivating mutations (exon vs. gene loss) (fig. S11B).

Since we do not consider frame-preserving deletions as inactivating mutations anymore, we added a new step to re-classify likely non-functional genes where most parts are lost due to frame-preserving deletions. To this end, we compute the percentage of reference codons that align to sense codons in the query (fig. S13) and classify a transcript as “uncertain loss” if this percentage is less than 50% and as “lost” if this percentage is less than 35%. Please note that by definition this percentage is 0% in case a gene is entirely deleted and spanning orthologous chains exist.

### Orthology inference

#### Classifying genes based on the classification of all transcripts and all orthologous loci

In the previous steps, TOGA aligns and classifies transcripts in the query genome. For orthology inference, individual predicted transcripts need to be consolidated into predicted genes. Importantly, a gene in the reference can have several transcripts (isoforms) and a given gene can have several inferred orthologous loci in the query. In the third step, TOGA uses all orthologous loci and the classification of all transcripts to determine whether the gene has at least one functional ortholog in the query and, if so, what the orthology type is (1:1, 1:many, many:1, many:many).

While transcripts in the reference are already assigned to genes in the input gene annotation, transcripts in the query need to be assigned to predicted genes. To this end, TOGA assigns two transcripts to the same gene, if their coding exons overlap by at least one base on the same strand (fig. S24A). This allows TOGA to correctly annotate and distinguish nested genes on the same strand and overlapping genes located in antisense orientation (fig. S24B and C).

For a given reference gene and one orthologous query locus, TOGA considers the classification of all transcripts of this gene that were annotated for this locus and applies the following order of precedence: “intact”, “partially intact”, “uncertain loss”, “lost”, “missing”, “paralogous projection” (fig. S11B). Hence, if at least one transcript is classified as intact, then TOGA infers that this orthologous locus contains a functional gene ortholog. An orthologous locus is inferred to contain a lost gene, if and only if all annotated transcripts of the given gene are classified as lost. To determine orthology type, TOGA then considers for each reference gene the classification of all its orthologous loci and for each of these query loci which reference genes were annotated.

#### Resolving many:many relationships supported by weak orthology

In the last step, TOGA uses the chain orthology probabilities computed by the gradient boosting approach (scores) to remove individual orthology relationships within a set of many:many orthologous genes that have substantially weaker support. For genes with a putative many:many orthology relationship, where ‘cross-gene’ orthology is supported only by alignment chains with weak orthology scores, this procedure aims at revealing the correct 1:1 orthology relationships. To this end, TOGA builds a bipartite graph with nodes representing reference and query genes and edges representing inferred orthology relationships weighted by the orthology score of the respective chain (fig. S25A). TOGA then tests if edges with substantially weaker orthology scores can be removed from the many:many orthology graph. To this end, TOGA subdivides all edges into two sets: set 1 contains all edges that connect a leaf node (reference or query gene that has only one inferred ortholog), set 2 contains all other edges. Let S_min_ be the minimum orthology score of edges in set 1. Branches in set 2 with a score < S_min_ * 0.9 will be removed (fig. S25B), unless one of the following conditions is true. First, no edge will be removed in the graph, if this would result in an isolated node that loses all its orthology connections (fig. S25D). Second, if two reference genes (say A and B) have more than one mutual orthology connection, TOGA does not remove edges that result in separating A and B into different orthology groups (fig. S25C). Third, in a complete bipartite graph, where every reference gene is connected to every query gene, no edge will be removed as there is no leaf in the graph (fig. S25E).

### Genome browser visualization

To visualize the annotations and gene/transcript classifications generated by TOGA in a genome browser, we extended the UCSC genome browser source code by a new TOGA annotation track type. The query annotations are loaded as a standard browser track in bed12 format and clicking on a transcript provides the following information: (i) the reference transcript with a link to Ensembl (or another user-defined gene resource) and reference genome coordinates, (ii) the orthology score of the chain used for projecting this transcript to the current locus, together with the features used for the machine learning classification, (iii) the transcript classification (intact, partial intact, etc.) together with the features that underlie this classification, (iv) a figure that visualizes all exons together with their class (present, missing, deleted) and all inactivating mutations, (v) a list of all detected inactivating mutations, (vi) the sequence alignment of the reference and the predicted query protein, and (vii) nucleotide alignments of individual exons together with coordinates, expected regions, %nucleotide identity and %BLOSUM values (fig. S42). This implementation comprises a new handler function in UCSC’s hgc.c that determines whether the user clicked on a TOGA annotation track and if so fetches all data from three SQL tables that hold the information described above. Instead of storing an exon visualization figure file for each transcript, we generate this visualization by including pre-computed SVG image code that is stored in the SQL table in the generated html page. The code additions to UCSC’s kent source are available on the TOGA github page in subdirectory ucsc_browser_visualization. Our UCSC browser mirror at https://genome.senckenberg.de/ provides the TOGA track functionality for 488 placental mammal, 21 non-placental mammal and 501 bird assemblies. Work is in progress to integrate TOGA tracks into UCSC’s GenArk by storing the TOGA data in bigBed files instead of SQL tables.

### Computing alignment chains

All 509 placental mammal alignment chains with human (hg38) and with mouse (mm10) as the reference were computed with the same parameters that are sufficiently sensitive to align orthologous exons between placental mammals (*48*). Briefly, we used LASTZ (version 1.04.00 or 1.04.03) (*49*) (parameters K = 2400, L = 3000, H = 2000, Y = 9400, default lastz scoring matrix) to generate local alignments. These local alignments were ‘chained’ using axtChain (*16*) (all parameters default except setting linearGap=loose). Next, we applied RepeatFiller (*50*) (all parameters default) to add previously missed alignments between repetitive regions and chainCleaner (*51*) (all parameters default except setting minBrokenChainScore = 75000 and specifying -doPairs) to improve alignment specificity. All 501 bird alignment chains with chicken (galGal6) as the reference and all chains with other reference species were computed in the same way.

We also compared TOGA using alignment chains that were generated by the UCSC genome browser group with less sensitive parameters and without RepeatFiller and chainCleaner. In these tests, we used human (hg38) as the reference and mouse (mm10), cow (bosTau9) and dog (canFam3) as three query species. As shown in fig. S45, with the sensitive alignment chains TOGA annotated 223, 120, and 114 additional orthologous genes for mouse, cow and dog, respectively, despite using the same query assemblies. This suggests that higher alignment sensitivity, obtained by different lastz parameter settings and the application of RepeatFiller and chainCleaner, makes it easier for TOGA to detect and annotate orthologs. Therefore, we recommend this workflow to generate chains for new assemblies.

To facilitate running the complex chain-generating procedure, we provide a pipeline that uses modified UCSC source code scripts and nextflow to execute the compute cluster-dependent steps. This pipeline was tested on different Linux systems and is available at https://github.com/hillerlab/make_lastz_chains.

### Application of TOGA

To use TOGA to infer orthologs and annotate genes in numerous mammalian genomes, we used the human GENCODE V38 (Ensembl 104) and the mouse GENCODE VM25 (Ensembl 100) gene annotation as reference. First, we extracted all transcripts for human and mouse from the Ensembl Biomart database (*22, 44*). In addition, we downloaded principal isoforms from the APPRIS database (*52*). Ideally, the input set of transcripts should be as comprehensive as possible to enable TOGA to also annotate alternative exons and splice sites; however, including problematic transcripts such as fusion transcripts or potential NMD targets can lead to wrong gene classifications or orthology types. Therefore, TOGA provides a script to filter the input set of transcripts as follows. First, all non-coding transcripts that lack an annotated CDS are excluded. Second, we excluded isoforms whose CDS is too short. To this end, we compute for each gene a CDS length threshold as 80% of the CDS length of the principal APPRIS isoform. If the gene has more than one principal isoform, we used the principal isoform with the shortest CDS. If APPRIS does not provide a principal isoform for the gene, we used the transcript with the longest CDS instead. We then excluded all transcripts that have CDS length below this threshold. Third, we excluded erroneous transcripts that have a CDS length not divisible by 3. Fourth, we excluded potential NMD targets that have the annotated stop codon more than 55 bp upstream of the last exon-exon junction. Fifth, we excluded isoforms that have introns shorter than 20 bp, as such micro-introns are often used to mask frameshifting mutations. Sixth, if several isoforms have an identical coding region, we selected only the one with the longest UTR. This step reduces redundancy as TOGA only annotates the CDS. Seventh, we excluded transcripts that have in-frame stop codons, unless the stop codon(s) is a TGA codon, in which case it may encode selenocysteine. Finally, we excluded transcripts that do not start with an ATG codon or end with a stop codon.

For genes with many transcripts, these filters ensure that only proper transcripts will be used as input for TOGA. However, it is possible that these filters eliminate all transcripts of a gene, for example, if the reference genome has a base error in a constitutive exon. Since this would result in missing the gene entirely, we include for such genes the longest transcript that has a CDS length divisible by three.

To apply TOGA with other mammals as reference, we obtained transcripts from the UCSC table ncbiRefSeq, holding the NCBI Felis catus Annotation Release 104 (2019-12-10) for cat (felCat9 assembly), NCBI Bos taurus Annotation Release 106 (2019-12-18) for cow (bosTau9) and NCBI Equus caballus Annotation Release 103 (2019-12-10) for horse (equCab3). To apply TOGA to birds, we used chicken (galGal6 assembly, NCBI accession GCA_000002315.5) as the reference. We downloaded the NCBI RefSeq annotation (GCF_000002315.6_GRCg6a_genomic.gff.gz) and combined this with the chicken APPRIS principal isoforms. To apply TOGA to other species, we downloaded NCBI RefSeq annotations (*21*) for the green sea turtle (GCF_015237465.1_rCheMyd1.pri_genomic.gff.gz), red-eared slider turtle (GCF_013100865.1_CAS_Tse_1.0_genomic.gff.gz), perch pike (GCF_008315115.2_SLUC_FBN_1.2_genomic.gff.gz), purple sea urchin (GCF_000002235.5_Spur_5.0_genomic.gff.gz), tobacco hawkmoth (GCF_014839805.1_JHU_Msex_v1.0_genomic.gff.gz), and *Arabidopsis thaliana* (GCF_000001735.4_TAIR10.1_genomic.gff.gz). These transcript sets were filtered as described above. For all non-mammalian genomes, we applied the standard TOGA method with default parameters and the machine learning model trained on human-mouse orthologs.

The assemblies of human (hg38) and mouse (mm10) also contain alternative haplotypes and structural variants (e.g. chr22_KI270876v1_alt). In case a haplotype contains the same gene as a reference chromosome (e.g. chr22), TOGA will infer an incorrect 2:1 orthology relationship in a query, since the reference gene is contained twice in the input annotation (at different genomic loci). To avoid this, we only considered for human chr1-chr22 and chrX and for mouse chr1-chr19 and chrX. Progress in sequencing and assembly allows it now to fully assemble both haplotypes of a diploid organism. For such assemblies, we recommend generating alignments and running TOGA individually on both haplotype assemblies, as recently demonstrated for the common vampire bat (*37*).

The final input annotations that TOGA used with human as the reference comprised 39,664 transcripts of 19,464 genes. For mouse, input annotations comprised 33,460 transcripts of 22,257 genes, and for chicken 38,252 transcripts of 18,039 genes.

Even for highly fragmented genome assemblies, low-scoring chains are extremely unlikely to represent orthologous parts of genes. Therefore, we did not classify chains with alignment scores <15,000 (a user adjustable threshold). To avoid excessive runtimes, we considered for each gene only the 100 highest-scoring orthologous chains in case the gene has more than 100 orthologous chains (such genes are part of larger gene families with many:many orthology relationships). To reduce runtime, we also considered genes as deleted, if the query locus defined by the closest up- and downstream alignment block is less than 5% of the total length of the reference CDS.

To count the number of annotated orthologs in a query species in Figure 5, we only considered genes that are classified by TOGA as intact, partially intact or uncertain loss.

### Gene loss detection accuracy

To evaluate TOGA gene loss detection pipeline sensitivity, we extracted a large set of conserved genes as a benchmark (Table S4). To this end, we extracted human genes that are annotated by Ensembl version 101 (downloaded July 8th, 2020) as 1:1 orthologs between human and mouse (mm10 assembly), rat (rn6), cow (bosTau9), and dog (canFam3). We excluded genes for which all isoforms contain very short introns (<50bp) in any of the four considered query species. This filter is necessary because such introns usually mask assembly base errors (frameshifting or stop codon mutations) or real inactivating mutations in lost genes (fig. S27). This resulted in a set of 11,161 human genes that are most likely conserved. Therefore, we considered all genes that TOGA classified as lost as false positives.

### Comparing ortholog detection between TOGA and Ensembl

We downloaded orthologous genes from Ensembl Biomart (version 104, downloaded August 12^th^ 2021) for human-rat (rn6 assembly), human-cow (bosTau9 assembly) and human-elephant (loxAfr3 assembly) together with the orthology type. Since TOGA but not Ensembl distinguishes between 1:many (more than 1 ortholog in the query species) and many:1 (one ortholog in the query, but more than one in the reference), we updated those Ensembl 1:many types as many:1, for which the orthology group had >1 gene annotated in reference and exactly one gene annotated in the query. For human-rat, we extracted for each Ensembl ortholog the orthology confidence value, the alignment identity between the “target and query gene” and the alignment coverage value from Ensembl Biomart. For each human-rat ortholog annotated by TOGA, we extracted TOGA’s orthology probability for the orthologous chain and computed the alignment identity and coverage value. These data are plotted in Figure 2D.

For the analysis of gene families, we downloaded gene families from the HUGO Gene Nomenclature Committee (*53*) (http://ftp.ebi.ac.uk/pub/databases/genenames/hgnc/tsv/hgnc_complete_set.txt) and used the Ensembl gene ID (ENSG) to determine gene families that comprise at least 30 members. Subfamilies of Zinc fingers, Olfactory receptors, T cell receptor and Immunoglobulin loci, and histones were combined. For genes for which only TOGA identified an ortholog, we then used the Ensembl gene ID to determine how many of these genes belong to larger gene families.

### Running BUSCO on genomes and annotations

For all tests that included mammalian BUSCO, we used BUSCO version 5.2.2 (*23*) and the mammalia odb10 dataset (downloaded on June 3rd, 2021) comprising 9,226 genes. The BUSCO odb10 datasets used for non-mammalian clades are specified in Figure 5D. To assess completeness of mammalian genome assemblies, we ran BUSCO in genome mode with default parameters using MetaEuk (version 34c21f2bf34c76f852c0441a29b104e5017f2f6d). To test whether there is a significant correlation between the BUSCO completeness and TOGA’s percent intact ancestral genes, we used the function cor.test() implemented in R version 4.0.3 and a two-sided test.

To assess completeness of gene annotations, we ran BUSCO in protein mode with default parameters and provided the protein sequences in a multi-fasta file as input. In contrast to applying BUSCO to a genome assembly, where one expects to find each of the “Universal Single-Copy Orthologs” only once in the assembly, applying BUSCO to a gene annotation results in the detection of many duplicated genes, because comprehensive annotations frequently include more than one transcript (splice variant) per gene. This does not indicate a problem but rather a comprehensive transcript annotation. For gene annotations, we therefore only report the number of completely detected BUSCO genes.

### Comparing the completeness of TOGA, Ensembl and NCBI annotations

For NCBI, we downloaded the annotated RefSeq protein sequences from the ftp server (ftp://ftp.ncbi.nlm.nih.gov/genomes/refseq/vertebrate_mammalian/) (protein.faa.gz files) for 118 placental mammals. For Ensembl release 104, we downloaded all annotated proteins (pep.all.fa.gz files) from http://ftp.ensembl.org/pub/current_fasta/ for 70 placental mammals. For TOGA, we used all annotated proteins obtained with human or with mouse as the reference. In addition, we also pooled the two TOGA protein sets. We used the NCBI RefSeq identifier and the assembly name provided by Ensembl to assure that all comparisons between TOGA and NCBI or Ensembl were done for the same genome assembly. We then ran BUSCO with the mammalia odb10 dataset on these sets of proteins, as described above.

### Adding TOGA as gene annotation evidence

To test whether TOGA as additional gene evidence can improve annotation completeness, we repeated the gene annotation procedure used in Jebb *et al.* (*6*), once with and once without TOGA. Briefly, we used EVidenceModeler (v1.1.1) (*54*) to combine previously generated gene evidence into a consensus gene set. Gene evidence comprised (i) *ab initio* gene predictions generated by Augustus (v3.3.1) with a bat-specific Augustus model (*55*), (ii) comparative gene predictions generated by Augustus CGP with a multiple genome alignment, (iii) full-length transcripts obtained from Iso-seq and RNA-seq data, and (iv) aligned protein and cDNA sequences of related bat species. These sources of evidence were weighted as in Jebb *et al.* (*6*) with *ab initio* predictions set to weight 1, comparative gene predictions and aligned proteins/cDNA sequences set to weight 2, RNA-seq transcripts set to weight 10, and Iso-seq transcripts set to weight 12. For the “with TOGA” annotation test, we used TOGA with human (hg38) as the reference and added transcripts classified as intact, partially intact or uncertain loss as an additional gene evidence with weight 8. We then used EVidenceModeler to split the genome into 1 Mb chunks with 150 kb overlap, determined consensus gene models and combined them into a genome-wide set. Afterwards, we added RNA- and Iso-seq transcripts that are not classified as NMD targets to the consensus transcript set. For the annotation that uses TOGA as an additional gene evidence, we also added TOGA-annotated transcripts classified as intact, partially intact or uncertain loss to the final transcript set. This resulted in two gene annotations for each of the six bats, one with and one without TOGA. Both annotations were assessed for completeness by applying BUSCO with the mammalian odb10 gene set to the annotated protein sequences.

We also tested the impact of adding aligned human proteins in addition to aligned proteins from closely related bats for two bats (*Myotis myotis* and *Rhinolophus ferrumequinum*). To this end, we downloaded the human reference proteome from https://ftp.uniprot.org/pub/databases/uniprot/current_release/knowledgebase/reference_proteomes/Eukaryota/UP000005640/UP000005640_9606.fasta.gz, which provides a BUSCO completeness of 99.5%. We used GenomeThreader (*56*) with the sensitive default parameters to align these proteins to the genomes of both bats. The aligned proteins were added to the other gene evidence and EVidenceModeler was used to generate a consensus gene set.

### Joining split genes in fragmented assemblies

To evaluate TOGA’s gene joining procedure, we used the TOGA annotations (human as reference) generated for the sperm whale (*Physeter macrocephalus*) and its closest relative, the pygmy sperm whale (*Kogia breviceps*). We first obtained a set of ‘benchmark’ genes for the contiguous *Physeter* genome GCA_002837175.2 assembly (Table S1). To this end, we extracted the longest CDS transcript for all genes that are classified as an intact 1:1 ortholog, that are located on a single scaffold, and for which all human exons are annotated in *Physeter*. For each transcript in this set, we determined whether TOGA annotated an intact 1:1 ortholog in the highly-fragmented *Kogia* assembly. We then determined whether this ortholog is located on a single *Kogia* scaffold (thus requiring no joining, which serves as a positive control) or was joined by TOGA from 2, 3 or ≥4 orthologous fragments. As a negative control, we extracted paralogs (instead of orthologs) in *Kogia* that are located on a single scaffold and for which all exons are annotated. To this end, we intentionally used TOGA to annotate exons in paralogous loci, obtained via chains whose orthology probability is <0.5. We produced pairwise alignments between the *Physeter* and *Kogia* sequences using MUSCLE version 3.8.1551 with default parameters and computed the nucleotide sequence identity.

To evaluate how effective the gene joining procedure is, we applied TOGA to *Kogia* and other highly-fragmented genomes. For each split gene, where TOGA joined orthologous fragments, we determined the CDS length and compared this to the CDS length of the longest-CDS transcript of the human ortholog. If the joined gene has a CDS length equal to the full-length human ortholog, this percentage is 100%. For comparison, we determined the CDS length of the single largest genomic fragment. Only split genes are shown in Figure 4C, but Table S10 provides data for all genes.

### Generalized linear models

To investigate factors that influence the number of orthologs annotated by TOGA across placental mammals, we fitted Poisson and negative binomial generalized linear models (GLMs) with log link functions in R (https://www.R-project.org/, version 4.1.2) using the packages stats and MASS (version 7.3-54) (*57*), respectively. Given that the distribution of ortholog counts was negatively skewed, we first transformed it by subtracting each value from the maximum value across the dataset. We then specified the transformed variable as the response in the GLMs. For predictors, we used (i) the divergence time to human in millions of years (obtained from the median value listed in http://timetree.org/), (ii) the evolutionary distance to human (number of substitutions per neutral site), (iii) the natural logarithm of the contig N50 value (base pairs), and (iv) the natural logarithm of the scaffold N50 value (base pairs). We fitted models with all possible combinations of these predictors, as well as an empty (intercept-only) model. To account for the strong positive correlation between evolutionary distance and divergence time, we specified both variables not as separate but as interacting predictors in models that included both.

The best-fitting model, determined through model selection according to the Akaike Information Criterion (AIC; Table S12), was a negative binomial GLM that included all four predictors. The coefficients of this model had P-values < 0.05. The variance-function-based R^2^ value (*58*), which we calculated using the R package rsq (https://CRAN.R-project.org/package=rsq, version 2.2), was 11.2%. By varying one predictor at a time and keeping the remaining predictors fixed at their mean values (fig. S38), we found that the most influential variable was contig N50, whereas the least influential was scaffold N50. Examining the distribution of AIC values across candidate GLMs (Table S12) led to the same conclusion. Performing the same analysis after excluding Hominoidea (Apes) led to qualitatively identical results and only slightly different model coefficients, P-values, and R^2^ values, indicating that our results are not biased by species that are very closely related to the reference genome (human). We also repeated this analysis including not only placental mammals but also monotremes and marsupials (fig. S38, Table S12).

### Ancestral placental mammal genes

To use TOGA to assess mammalian genome completeness and quality, we obtained a set of protein-coding genes that likely already existed in the placental mammal ancestor. Given that the basal split of placental mammals is not yet resolved (*59*), we conservatively defined ancestral placental mammal genes as those that have an intact reading frame in representatives of all three superorders: Boreoeutheria, Afrotheria and Xenarthra. We used the human GENCODE V38 (Ensembl 104) gene annotation (*22*), which implies that each gene is intact in Boreoeutheria, and then selected those genes that are classified by TOGA as intact or partially intact in at least one afrotherian and at least one xenarthran genome. We considered 11 afrotherian species (dugong, manatee, Asiatic elephant, African savanna elephant, cape rock hyrax, yellow-spotted hyrax, aardvark, cape golden mole, Talazac’s shrew tenrec, small Madagascar hedgehog, cape elephant shrew) and five xenarthran species (Hoffmann’s two-fingered sloth, southern two-toed sloth, giant anteater, southern tamandua, nine-banded armadillo). This procedure resulted in 18,430 genes (Table S13).

## Acknowledgment

We thank the UCSC genome browser group for providing software, Ensembl and NCBI for annotations, Ingo Ebersberger and Kerstin Lindblad-Toh for helpful comments, Franziska Friedrich for the TOGA logo, and the Computer Service Facilities of the MPI-CBG and MPI-PKS and Christoph Sinai for technical support.

## Author contributions

BMK implemented TOGA, generated and analyzed data. CM, EO, DJ, VS, MB, AEM, AWA, DGK, LH and MH contributed to data analysis. MH conceived and supervised the study, wrote the manuscript and made figures with help from BMK and other authors. All authors approved the final manuscript.

## Data and materials availability

The TOGA source code used for this study, and all scripts to run TOGA, create training and test data sets and browser tracks are permanently archived at Zenodo (*60*). Further code development will be tracked on https://github.com/hillerlab/TOGA. We recommend generating alignment chains with our pipeline (https://github.com/hillerlab/make_lastz_chains). All data are available in the manuscript or the supplementary material, and available for download at http://genome.senckenberg.de/download/TOGA/ and for browsing in our UCSC genome browser mirror at https://genome.senckenberg.de.

## Zoonomia Consortium

Gregory Andrews^1^, Joel C. Armstrong^2^, Matteo Bianchi^3^, Bruce W. Birren^4^, Kevin R. Bredemeyer^5^, Ana M. Breit^6^, Matthew J. Christmas^3^, Hiram Clawson^2^, Joana Damas^7^, Federica Di Palma^8,9^, Mark Diekhans^2^, Michael X. Dong^3^, Eduardo Eizirik^10^, Kaili Fan^1^, Cornelia Fanter^11^, Nicole M. Foley^5^, Karin Forsberg-Nilsson^12,13^, Carlos J. Garcia^14^, John Gatesy^15^, Steven Gazal^16^, Diane P. Genereux^4^, Linda Goodman^17^, Jenna Grimshaw^14^, Michaela K. Halsey^14^, Andrew J. Harris^5^, Glenn Hickey^18^, Michael Hiller^19,20,21^, Allyson G. Hindle^11^, Robert M. Hubley^22^, Graham M. Hughes^23^, Jeremy Johnson^4^, David Juan^24^, Irene M. Kaplow^25,26^, Elinor K. Karlsson^1,4,27^, Kathleen C. Keough^17,28,29^, Bogdan Kirilenko^19,20,21^, Klaus-Peter Koepfli^30,31,32^, Jennifer M. Korstian^14^, Amanda Kowalczyk^25,26^, Sergey V. Kozyrev^3^, Alyssa J. Lawler^4,26,33^, Colleen Lawless^23^, Thomas Lehmann^34^, Danielle L. Levesque^6^, Harris A. Lewin^7,35,36^, Xue Li^1,4,37^, Abigail Lind^28,29^, Kerstin Lindblad-Toh^3,4^, Ava Mackay-Smith^38^, Voichita D. Marinescu^3^, Tomas Marques-Bonet^39,40,41,42^, Victor C. Mason^43^, Jennifer R. S. Meadows^3^, Wynn K. Meyer^44^, Jill E. Moore^1^, Lucas R. Moreira^1,4^, Diana D. Moreno-Santillan^14^, Kathleen M. Morrill^1,4,37^, Gerard Muntané^24^, William J. Murphy^5^, Arcadi Navarro^39,41,45,46^, Martin Nweeia^47,48,49,50^, Sylvia Ortmann^51^, Austin Osmanski^14^, Benedict Paten^2^, Nicole S. Paulat^14^, Andreas R. Pfenning^25,26^, BaDoi N. Phan^25,26,52^, Katherine S. Pollard^28,29,53^, Henry E. Pratt^1^, David A. Ray^14^, Steven K. Reilly^38^, Jeb R. Rosen^22^, Irina Ruf^54^, Louise Ryan^23^, Oliver A. Ryder^55,56^, Pardis C. Sabeti^4,57,58^, Daniel E. Schäffer^25^, Aitor Serres^24^, Beth Shapiro^59,60^, Arian F. A. Smit^22^, Mark Springer^61^, Chaitanya Srinivasan^25^, Cynthia Steiner^55^, Jessica M. Storer^22^, Kevin A. M. Sullivan^14^, Patrick F. Sullivan^62,63^, Elisabeth Sundström^3^, Megan A. Supple^59^, Ross Swofford^4^, Joy-El Talbot^64^, Emma Teeling^23^, Jason Turner-Maier^4^, Alejandro Valenzuela^24^, Franziska Wagner^65^, Ola Wallerman^3^, Chao Wang^3^, Juehan Wang^16^, Zhiping Weng^1^, Aryn P. Wilder^55^, Morgan E. Wirthlin^25,26,66^, James R. Xue^4,57^, Xiaomeng Zhang^4,25,26^

Affiliations:

^1^Program in Bioinformatics and Integrative Biology, UMass Chan Medical School; Worcester, MA 01605, USA.

^2^Genomics Institute, University of California Santa Cruz; Santa Cruz, CA 95064, USA.

^3^Department of Medical Biochemistry and Microbiology, Science for Life Laboratory, Uppsala University; Uppsala, 751 32, Sweden.

^4^Broad Institute of MIT and Harvard; Cambridge, MA 02139, USA.

^5^Veterinary Integrative Biosciences, Texas A&M University; College Station, TX 77843, USA.

^6^School of Biology and Ecology, University of Maine; Orono, ME 04469, USA.

^7^The Genome Center, University of California Davis; Davis, CA 95616, USA.

^8^Genome British Columbia; Vancouver, BC, Canada.

^9^School of Biological Sciences, University of East Anglia; Norwich, UK.

^10^School of Health and Life Sciences, Pontifical Catholic University of Rio Grande do Sul; Porto Alegre, 90619-900, Brazil.

^11^School of Life Sciences, University of Nevada Las Vegas; Las Vegas, NV 89154, USA.

^12^Biodiscovery Institute, University of Nottingham; Nottingham, UK.

^13^Department of Immunology, Genetics and Pathology, Science for Life Laboratory, Uppsala University; Uppsala, 751 85, Sweden.

^14^Department of Biological Sciences, Texas Tech University; Lubbock, TX 79409, USA.

^15^Division of Vertebrate Zoology, American Museum of Natural History; New York, NY 10024, USA.

^16^Keck School of Medicine, University of Southern California; Los Angeles, CA 90033, USA.

^17^Fauna Bio Incorporated; Emeryville, CA 94608, USA.

^18^Baskin School of Engineering, University of California Santa Cruz; Santa Cruz, CA 95064, USA.

^19^Faculty of Biosciences, Goethe-University; 60438 Frankfurt, Germany.

^20^LOEWE Centre for Translational Biodiversity Genomics; 60325 Frankfurt, Germany.

^21^Senckenberg Research Institute; 60325 Frankfurt, Germany.

^22^Institute for Systems Biology; Seattle, WA 98109, USA.

^23^School of Biology and Environmental Science, University College Dublin; Belfield, Dublin 4, Ireland.

^24^Department of Experimental and Health Sciences, Institute of Evolutionary Biology (UPF-CSIC), Universitat Pompeu Fabra; Barcelona, 08003, Spain.

^25^Department of Computational Biology, School of Computer Science, Carnegie Mellon University; Pittsburgh, PA 15213, USA.

^26^Neuroscience Institute, Carnegie Mellon University; Pittsburgh, PA 15213, USA.

^27^Program in Molecular Medicine, UMass Chan Medical School; Worcester, MA 01605, USA.

^28^Department of Epidemiology & Biostatistics, University of California San Francisco; San Francisco, CA 94158, USA.

^29^Gladstone Institutes; San Francisco, CA 94158, USA.

^30^Center for Species Survival, Smithsonian’s National Zoo and Conservation Biology Institute; Washington, DC 20008, USA.

^31^Computer Technologies Laboratory, ITMO University; St. Petersburg 197101, Russia.

^32^Smithsonian-Mason School of Conservation, George Mason University; Front Royal, VA 22630, USA.

^33^Department of Biological Sciences, Mellon College of Science, Carnegie Mellon University; Pittsburgh, PA 15213, USA.

^34^Senckenberg Research Institute and Natural History Museum Frankfurt; 60325 Frankfurt am Main, Germany.

^35^Department of Evolution and Ecology, University of California Davis; Davis, CA 95616, USA.

^36^John Muir Institute for the Environment, University of California Davis; Davis, CA 95616, USA.

^37^Morningside Graduate School of Biomedical Sciences, UMass Chan Medical School; Worcester, MA 01605, USA.

^38^Department of Genetics, Yale School of Medicine; New Haven, CT 06510, USA.

^39^Catalan Institution of Research and Advanced Studies (ICREA); Barcelona, 08010, Spain.

^40^CNAG-CRG, Centre for Genomic Regulation, Barcelona Institute of Science and Technology (BIST); Barcelona, 08036, Spain.

^41^Department of Medicine and LIfe Sciences, Institute of Evolutionary Biology (UPF-CSIC), Universitat Pompeu Fabra; Barcelona, 08003, Spain.

^42^Institut Català de Paleontologia Miquel Crusafont, Universitat Autònoma de Barcelona; 08193, Cerdanyola del Vallès, Barcelona, Spain.

^43^Institute of Cell Biology, University of Bern; 3012, Bern, Switzerland.

^44^Department of Biological Sciences, Lehigh University; Bethlehem, PA 18015, USA.

^45^BarcelonaBeta Brain Research Center, Pasqual Maragall Foundation; Barcelona, 08005, Spain.

^46^CRG, Centre for Genomic Regulation, Barcelona Institute of Science and Technology (BIST); Barcelona, 08003, Spain.

^47^Department of Comprehensive Care, School of Dental Medicine, Case Western Reserve University; Cleveland, OH 44106, USA.

^48^Department of Vertebrate Zoology, Canadian Museum of Nature; Ottawa, Ontario K2P 2R1, Canada.

^49^Department of Vertebrate Zoology, Smithsonian Institution; Washington, DC 20002, USA.

^50^Narwhal Genome Initiative, Department of Restorative Dentistry and Biomaterials Sciences, Harvard School of Dental Medicine; Boston, MA 02115, USA.

^51^Department of Evolutionary Ecology, Leibniz Institute for Zoo and Wildlife Research; 10315 Berlin, Germany.

^52^Medical Scientist Training Program, University of Pittsburgh School of Medicine; Pittsburgh, PA 15261, USA.

^53^Chan Zuckerberg Biohub; San Francisco, CA 94158, USA

^54^Division of Messel Research and Mammalogy, Senckenberg Research Institute and Natural History Museum Frankfurt; 60325 Frankfurt am Main, Germany.

^55^Conservation Genetics, San Diego Zoo Wildlife Alliance; Escondido, CA 92027, USA.

^56^Department of Evolution, Behavior and Ecology, School of Biological Sciences, University of California San Diego; La Jolla, CA 92039, USA.

^57^Department of Organismic and Evolutionary Biology, Harvard University; Cambridge, MA 02138, USA.

^58^Howard Hughes Medical Institute; Chevy Chase, MD, USA.

^59^Department of Ecology and Evolutionary Biology, University of California Santa Cruz; Santa Cruz, CA 95064, USA.

^60^Howard Hughes Medical Institute, University of California Santa Cruz; Santa Cruz, CA 95064, USA.

^61^Department of Evolution, Ecology and Organismal Biology, University of California Riverside; Riverside, CA 92521, USA.

^62^Department of Genetics, University of North Carolina Medical School; Chapel Hill, NC 27599, USA.

^63^Department of Medical Epidemiology and Biostatistics, Karolinska Institutet; Stockholm, Sweden.

^64^Iris Data Solutions, LLC; Orono, ME 04473, USA.

^65^Museum of Zoology, Senckenberg Natural History Collections Dresden; 01109 Dresden, Germany.

^66^Allen Institute for Brain Science; Seattle, WA 98109, USA

## Supplementary Materials

Tables S1 – S15

Figs S1 – S45

References (*41-59, 61-68*)

## References and Notes

1. T. Gabaldon, E. V. Koonin, Functional and evolutionary implications of gene orthology. Nature reviews. Genetics 14, 360–366 (2013).

2. P. Kapli, Z. Yang, M. J. Telford, Phylogenetic tree building in the genomic age. Nature reviews. Genetics 21, 428–444 (2020).

3. A. M. Altenhoff, R. A. Studer, M. Robinson-Rechavi, C. Dessimoz, Resolving the ortholog conjecture: orthologs tend to be weakly, but significantly, more similar in function than paralogs. PLoS computational biology 8, e1002514 (2012).

4. J. Huerta-Cepas et al., Fast Genome-Wide Functional Annotation through Orthology Assignment by eggNOG-Mapper. Molecular biology and evolution 34, 2115–2122 (2017).

5. V. Sharma et al., A genomics approach reveals insights into the importance of gene losses for mammalian adaptations. Nature communications 9, 1215 (2018).

6. D. Jebb et al., Six reference-quality genomes reveal evolution of bat adaptations. Nature 583, 578–584 (2020).

7. A. M. Altenhoff, N. M. Glover, C. Dessimoz, Inferring Orthology and Paralogy. Methods in molecular biology 1910, 149–175 (2019).

8. L. Li, C. J. Stoeckert, Jr., D. S. Roos, OrthoMCL: identification of ortholog groups for eukaryotic genomes. Genome Res 13, 2178–2189 (2003).

9. C. M. Train, N. M. Glover, G. H. Gonnet, A. M. Altenhoff, C. Dessimoz, Orthologous Matrix (OMA) algorithm 2.0: more robust to asymmetric evolutionary rates and more scalable hierarchical orthologous group inference. Bioinformatics 33, i75–i82 (2017).

10. E. M. Zdobnov et al., OrthoDB v9.1: cataloging evolutionary and functional annotations for animal, fungal, plant, archaeal, bacterial and viral orthologs. Nucleic Acids Res 45, D744–D749 (2017).

11. L. J. Jensen et al., eggNOG: automated construction and annotation of orthologous groups of genes. Nucleic Acids Res 36, D250–254 (2008).

12. D. M. Emms, S. Kelly, OrthoFinder: phylogenetic orthology inference for comparative genomics. Genome Biol 20, 238 (2019).

13. J. Huerta-Cepas, S. Capella-Gutierrez, L. P. Pryszcz, M. Marcet-Houben, T. Gabaldon, PhylomeDB v4: zooming into the plurality of evolutionary histories of a genome. Nucleic Acids Res 42, D897–902 (2014).

14. A. J. Vilella et al., EnsemblCompara GeneTrees: Complete, duplication-aware phylogenetic trees in vertebrates. Genome Res 19, 327–335 (2009).

15. K. Trachana et al., Orthology prediction methods: a quality assessment using curated protein families. Bioessays 33, 769–780 (2011).

16. W. J. Kent, R. Baertsch, A. Hinrichs, W. Miller, D. Haussler, Evolution’s cauldron: duplication, deletion, and rearrangement in the mouse and human genomes. Proceedings of the National Academy of Sciences of the United States of America 100, 11484–11489 (2003).

17. R. P. Meisel, T. Connallon, The faster-X effect: integrating theory and data. Trends in Genetics 29, 537–544 (2013).

18. V. Sharma, P. Schwede, M. Hiller, CESAR 2.0 substantially improves speed and accuracy of comparative gene annotation. Bioinformatics 33, 3985–3987 (2017).

19. V. Sharma, A. Elghafari, M. Hiller, Coding exon-structure aware realigner (CESAR) utilizes genome alignments for accurate comparative gene annotation. Nucleic Acids Res 44, e103 (2016).

20. F. Thibaud-Nissen, et al., P8008 The NCBI Eukaryotic Genome Annotation Pipeline. Journal of Animal Science 94, 184–184 (2016).

21. N. A. O’Leary et al., Reference sequence (RefSeq) database at NCBI: current status, taxonomic expansion, and functional annotation. Nucleic Acids Res 44, D733–745 (2016).

22. A. Frankish, et al., Gencode 2021. Nucleic Acids Res 49, D916–D923 (2020).

23. M. Manni, M. R. Berkeley, M. Seppey, F. A. Simao, E. M. Zdobnov, BUSCO Update: Novel and Streamlined Workflows along with Broader and Deeper Phylogenetic Coverage for Scoring of Eukaryotic, Prokaryotic, and Viral Genomes. Molecular biology and evolution 38, 4647–4654 (2021).

24. M. Stanke, O. Schoffmann, B. Morgenstern, S. Waack, Gene prediction in eukaryotes with a generalized hidden Markov model that uses hints from external sources. BMC Bioinformatics 7, 62 (2006).

25. S. Konig, L. W. Romoth, L. Gerischer, M. Stanke, Simultaneous gene finding in multiple genomes. Bioinformatics 32, 3388–3395 (2016).

26. A. Rhie et al., Towards complete and error-free genome assemblies of all vertebrate species. Nature 592, 737–746 (2021).

27. Zoonomia Consortium, A comparative genomics multitool for scientific discovery and conservation. Nature 587, 240–245 (2020).

28. S. Feng et al., Dense sampling of bird diversity increases power of comparative genomics. Nature 587, 252–257 (2020).

29. G. Fan et al., The first chromosome-level genome for a marine mammal as a resource to study ecology and evolution. Mol Ecol Resour 19, 944–956 (2019).

30. F. S. Sharko et al., Steller’s sea cow genome suggests this species began going extinct before the arrival of Paleolithic humans. Nature communications 12, 2215 (2021).

31. W. J. Murphy, N. M. Foley, K. R. Bredemeyer, J. Gatesy, M. S. Springer, Phylogenomics and the Genetic Architecture of the Placental Mammal Radiation. Annu Rev Anim Biosci 9, 29–53 (2020).

32. E. D. Jarvis et al., Whole-genome analyses resolve early branches in the tree of life of modern birds. Science 346, 1320–1331 (2014).

33. V. Ranwez, E. J. P. Douzery, C. Cambon, N. Chantret, F. Delsuc, MACSE v2: Toolkit for the Alignment of Coding Sequences Accounting for Frameshifts and Stop Codons. Molecular biology and evolution 35, 2582–2584 (2018).

34. B. T. Lee et al., The UCSC Genome Browser database: 2022 update. Nucleic Acids Res 50, D1115–D1122 (2021).

35. J. Damas et al., Broad host range of SARS-CoV-2 predicted by comparative and structural analysis of ACE2 in vertebrates. Proceedings of the National Academy of Sciences of the United States of America 117, 22311–22322 (2020).

36. J. G. Roscito et al., Convergent and lineage-specific genomic differences in limb regulatory elements in limbless reptile lineages. Cell Rep 38, 110280 (2022).

37. M. Blumer et al., Gene losses in the common vampire bat illuminate molecular adaptations to blood feeding. Sci Adv 8, eabm6494 (2022).

38. H. Indrischek et al., Vision-related convergent gene losses reveal SERPINE3’s unknown role in the eye. eLife 11, (2022).

39. H. A. Lewin et al., Earth BioGenome Project: Sequencing life for the future of life. Proceedings of the National Academy of Sciences of the United States of America 115, 4325–4333 (2018).

40. E. Teeling, et al., Bat Biology, Genomes, and the Bat1K Project: To Generate Chromosome-Level Genomes for all Living Bat Species. Annu Rev Anim Biosci 6, 23–46 (2017).

41. J. Lehmann, P. F. Stadler, S. J. Prohaska, SynBlast: assisting the analysis of conserved synteny information. BMC Bioinformatics 9, 351 (2008).

42. J. Jun, Mandoiu, II, C. E. Nelson, Identification of mammalian orthologs using local synteny. BMC Genomics 10, 630 (2009).

43. S. Jahangiri-Tazehkand, L. Wong, C. Eslahchi, OrthoGNC: A Software for Accurate Identification of Orthologs Based on Gene Neighborhood Conservation. Genomics Proteomics Bioinformatics 15, 361–370 (2017).

44. A. D. Yates, et al., Ensembl 2020. Nucleic Acids Res 48, D682–D688 (2019).

45. T. Chen, C. Guestrin, paper presented at the Proceedings of the 22nd ACM SIGKDD International Conference on Knowledge Discovery and Data Mining, San Francisco, California, USA, 2016.

46. A. Levine, R. Durbin, A computational scan for U12-dependent introns in the human genome sequence. Nucleic Acids Res 29, 4006–4013 (2001).

47. T. S. Alioto, U12DB: a database of orthologous U12-type spliceosomal introns. Nucleic Acids Res 35, D110–115 (2007).

48. V. Sharma, M. Hiller, Increased alignment sensitivity improves the usage of genome alignments for comparative gene annotation. Nucleic Acids Res 45, 8369–8377 (2017).

49. R. S. Harris, The Pennsylvania State University, (2007).

50. E. Osipova, N. Hecker, M. Hiller, RepeatFiller newly identifies megabases of aligning repetitive sequences and improves annotations of conserved non-exonic elements. Gigascience 8, giz132 (2019).

51. H. G. Suarez, B. E. Langer, P. Ladde, M. Hiller, chainCleaner improves genome alignment specificity and sensitivity. Bioinformatics 33, 1596–1603 (2017).

52. J. M. Rodriguez, et al., APPRIS 2017: principal isoforms for multiple gene sets. Nucleic Acids Res 46, D213–D217 (2017).

53. L. C. Daugherty, R. L. Seal, M. W. Wright, E. A. Bruford, Gene family matters: expanding the HGNC resource. Hum Genomics 6, 4 (2012).

54. B. J. Haas et al., Automated eukaryotic gene structure annotation using EVidenceModeler and the Program to Assemble Spliced Alignments. Genome Biol 9, R7 (2008).

55. M. Stanke, M. Diekhans, R. Baertsch, D. Haussler, Using native and syntenically mapped cDNA alignments to improve de novo gene finding. Bioinformatics 24, 637–644 (2008).

56. G. Gremme, V. Brendel, M. E. Sparks, S. Kurtz, Engineering a Software Tool for Gene Structure Prediction in Higher Organisms. Information and Software Technology 47, 965–978 (2005).

57. W. N. Venables, B. D. Ripley, Modern Applied Statistics with S. (Springer, 2002), vol. Fourth Edition.

58. D. Zhang, A Coefficient of Determination for Generalized Linear Models. The American Statistician 71, 310–316 (2017).

59. N. M. Foley, M. S. Springer, E. C. Teeling, Mammal madness: is the mammal tree of life not yet resolved? *Philosophical transactions of the Royal Society of London. Series B*, Biological sciences 371, 20150140 (2016).

60. B. M. Kirilenko, M. Hiller, TOGA source code v1.0.0. https://zenodo.org/record/6400671 (2022).

61. N. Hecker, V. Sharma, M. Hiller, Convergent gene losses illuminate metabolic and physiological changes in herbivores and carnivores. Proceedings of the National Academy of Sciences of the United States of America 116, 3036–3041 (2019).

62. M. Huelsmann et al., Genes lost during the transition from land to water in cetaceans highlight genomic changes associated with aquatic adaptations. Sci Adv 5, eaaw6671 (2019).

63. V. Sharma, N. Hecker, F. Walther, H. Stuckas, M. Hiller, Convergent Losses of TLR5 Suggest Altered Extracellular Flagellin Detection in Four Mammalian Lineages. Molecular biology and evolution 37, 1847–1854 (2020).

64. L. M. Boyden et al., Skint1, the prototype of a newly identified immunoglobulin superfamily gene cluster, positively selects epidermal gammadelta T cells. Nat Genet 40, 656–662 (2008).

65. T. Narita, T. Nitta, S. Nitta, T. Okamura, H. Takayanagi, Mice lacking all of the Skint family genes. Int Immunol 30, 301–309 (2018).

66. R. H. Mohamed et al., The SKINT1-like gene is inactivated in hominoids but not in all primate species: implications for the origin of dendritic epidermal T cells. PloS one 10, e0123258 (2015).

67. P. A. Morin et al., Reference genome and demographic history of the most endangered marine mammal, the vaquita. Molecular Ecology Resources 21, 1008–1020 (2021).

68. A. Loytynoja, Phylogeny-aware alignment with PRANK. Methods in molecular biology 1079, 155–170 (2014).

